# Impact of preanalytical factors on liquid biopsy in the canine cancer model

**DOI:** 10.1101/2024.07.29.605605

**Authors:** Kate Megquier, Christopher Husted, Justin Rhoades, Michelle E. White, Diane P. Genereux, Frances L. Chen, Kan Xiong, Euijin Kwon, Ross Swofford, Corrie Painter, Viktor Adalsteinsson, Cheryl A. London, Heather L. Gardner, Elinor K. Karlsson

**Affiliations:** Broad Institute of MIT and Harvard, Cambridge, MA, USA; Genomics and Computational Biology, UMass Chan Medical School, Worcester, MA, USA; Morningside Graduate School of Biomedical Sciences, UMass Chan Medical School, Worcester, MA, USA; Tufts Cummings School of Veterinary Medicine, North Grafton, MA, USA; Program in Molecular Medicine, UMass Chan Medical School, Worcester, MA, USA

**Author notes:** Correspondence: Elinor K. Karlsson, PhD UMass Chan Medical School, 368 Plantation Street, AS4.1065 Worcester MA 01605; Kate Megquier, DVM, PhD, Broad Institute of MIT and Harvard 415 Main Street, 3011B Cambridge, MA 02142. Authors contributed equally (Heather L. Gardner, Elinor K. Karlsson).

**Keywords:** liquid biopsy, cell-free DNA, cancer, comparative, canine, optimization

## Abstract

While liquid biopsy has potential to transform cancer diagnostics through minimally-invasive detection and monitoring of tumors, the impact of preanalytical factors such as the timing and anatomical location of blood draw is not well understood. To address this gap, we leveraged pet dogs with spontaneous cancer as a model system, as their compressed disease timeline facilitates rapid diagnostic benchmarking. Key liquid biopsy metrics from dogs were consistent with existing reports from human patients. The tumor content of samples was higher from venipuncture sites closer to the tumor and from a central vein. Metrics also differed between lymphoma and non-hematopoietic cancers, urging cancer-type-specific interpretation. Liquid biopsy was highly sensitive to disease status, with changes identified soon after post chemotherapy administration, and trends of increased tumor fraction and other metrics observed prior to clinical relapse in dogs with lymphoma or osteosarcoma. These data support the utility of pet dogs with cancer as a relevant system for advancing liquid biopsy platforms.

## INTRODUCTION

Liquid biopsy is a powerful, minimally-invasive method to detect, monitor, and evaluate the genomic landscape of tumors. Cell-free DNA (cfDNA), including circulating tumor DNA derived from tumors, is collected from blood and sequenced, enabling comprehensive assessment of tumor burden and somatic mutations^1^. This technique has the potential to enable precision cancer treatment and monitoring by matching patients to targeted therapies^2–5^, identifying molecular relapse prior to clinical relapse^6–10^, and detecting emerging mutations that confer therapeutic resistance^11^.

While liquid biopsy methodologies have advanced substantially, it is still not entirely clear how a variety of preanalytical variables impact cfDNA yield and diagnostic results. These include patient-specific factors, cancer type, and sample collection protocols, among others^12^. The performance of liquid biopsy is reliant on collecting a sufficient amount of tumor DNA, which may make up only a small fraction of total cfDNA from a given blood sample. Increasing the yield of tumor DNA can therefore substantially impact test sensitivity and specificity^12,13^. Recent data has shown that cfDNA levels vary by collection tube^14^, cancer type^15^, by time of day^16,17^, and timing of physical exercise^18–21^. However, there are currently no standardized recommendations or guidance regarding sample collection to mitigate variability and optimize reliability, accuracy, and cost efficiency in the clinical setting^22,23^.

Discerning the influence of preanalytical factors on liquid biopsies is challenging in human studies as clinical protocols typically dictate that blood be drawn from the median cubital vein^24^. Moreover, it is often difficult to intensively sample human patients over short timelines and as such, they may be collected weeks, months, or even years apart^25^. While mouse models have been used to investigate liquid biopsy application, several constraints exist, including very small sample volumes, limited options for blood draw site, and challenges following individual mice over the course of diagnosis, remission and eventual therapeutic resistance^26–28^. Importantly, mouse cancer models also do not capture the environmental and lifestyle diversity of human cohorts^26^. Mouse models also tend to be more genetically homogeneous^26^.

Use of pet dogs with cancer as a relevant comparative and preclinical model has recently gained traction as a mechanism to more effectively bridge translation of findings in mice to human patients. Cancers in pet dogs develop spontaneously and possess similar histologies, clinical characteristics, and genomic alterations as human cancers^29^, and it is estimated that over 1.5 million dogs are diagnosed with cancer each year^30,31^. With their shorter lifespans and disease course, longitudinal clinical studies in dogs can be completed in 1-2 years, instead of the 5-10+ years necessary for similar human studies^29,32,33^. Dogs with cancer are treated with the same modalities used in human medicine, including chemotherapy, stereotactic/conformal radiation therapy, immunotherapy, and small molecule inhibitors^29^. However, veterinary clinical protocols are more amenable to modification and evaluation of novel approaches in the research setting^19^. Several studies have already established the feasibility of isolating cfDNA from dogs and analyzing it via PCR, targeted, or whole-genome sequencing NGS-based approaches to identify tumor-derived mutations^34–49^.

Here, we demonstrate the utility of the comparative canine model for optimizing liquid biopsy techniques. We first credentialled the model by confirming concordance of somatic mutations between tumor and paired cfDNA. We then assessed the effect of patient characteristics and preanalytical variables such as blood draw site and time of day, and investigated cfDNA dynamics in relationship to clinical course and outcomes over short and long time scales (Fig 1), confirming the power of pet dogs as a cancer model.

**Figure 1.**
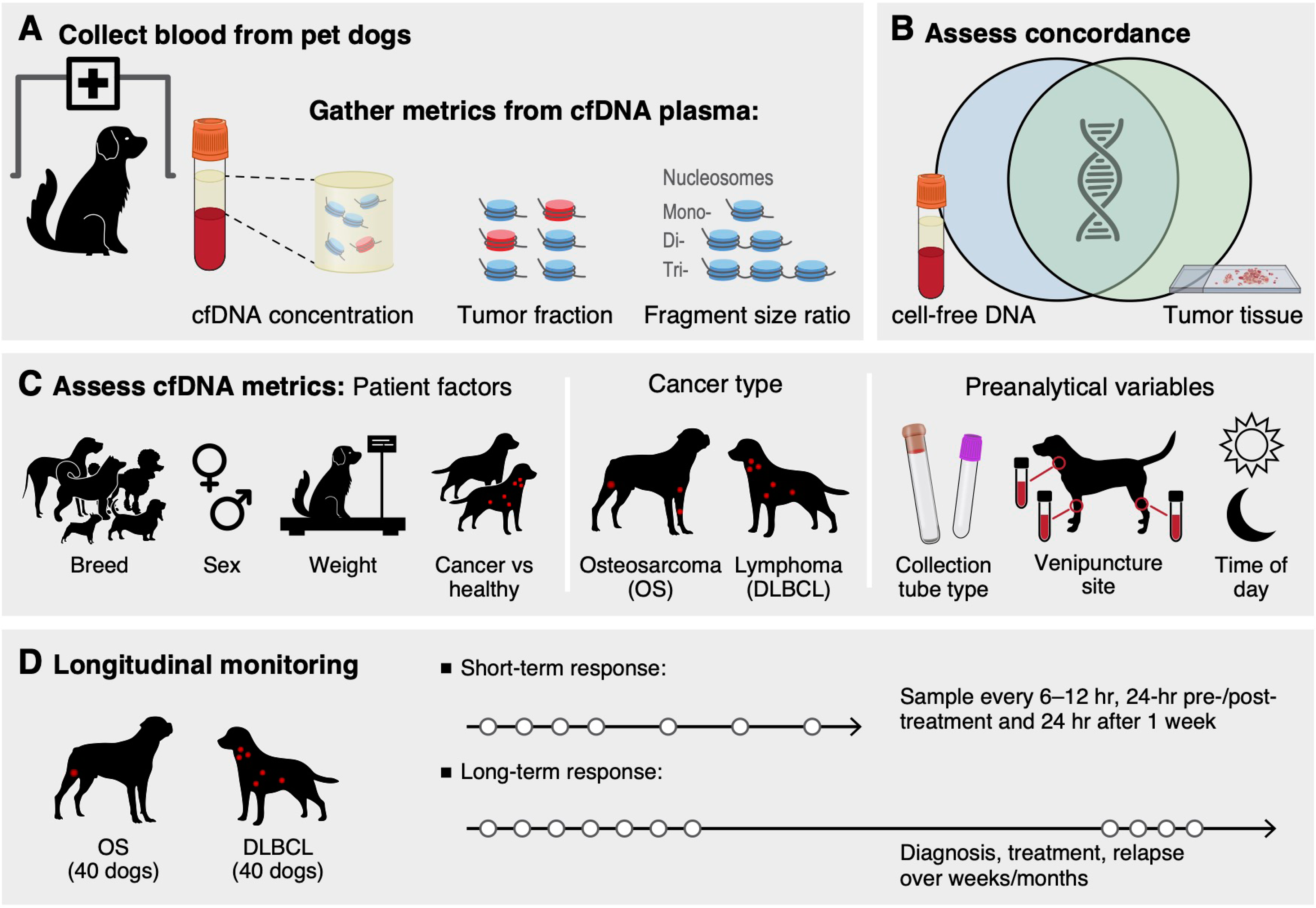
Overview. Illustration of the study aims and methods.

## RESULTS

### Enrollment and sample collection

We evaluated cfDNA samples from a diverse set of 149 dogs—133 with cancer and 16 controls without a cancer diagnosis. The number of samples collected from each dog varied from 1 to 11 depending on the study cohort (Fig 2A), totaling 430 individual cfDNA samples. Overall, the dogs in our sample set were representative of the U.S. dog population (Table S1). Dogs with cancer were not statistically different from the healthy dogs in terms of sex (p_fisher_=0.6), spay/neuter status (p_fisher_=0.058; only 6.5% of dogs were intact), weight (p_t-test_=0.63) or whether they had ancestry from one or multiple breeds (p_fisher_=0.17) (Fig SDOGS A-D). Dogs with cancer were older (8.5+/-2.7 years old for 111 dogs with cancer and 4.4+/-3.2 years old for 16 healthy dogs, p_t-test_=1.2x10^-4^) (Fig SDOGS E).

**Figure 2.**
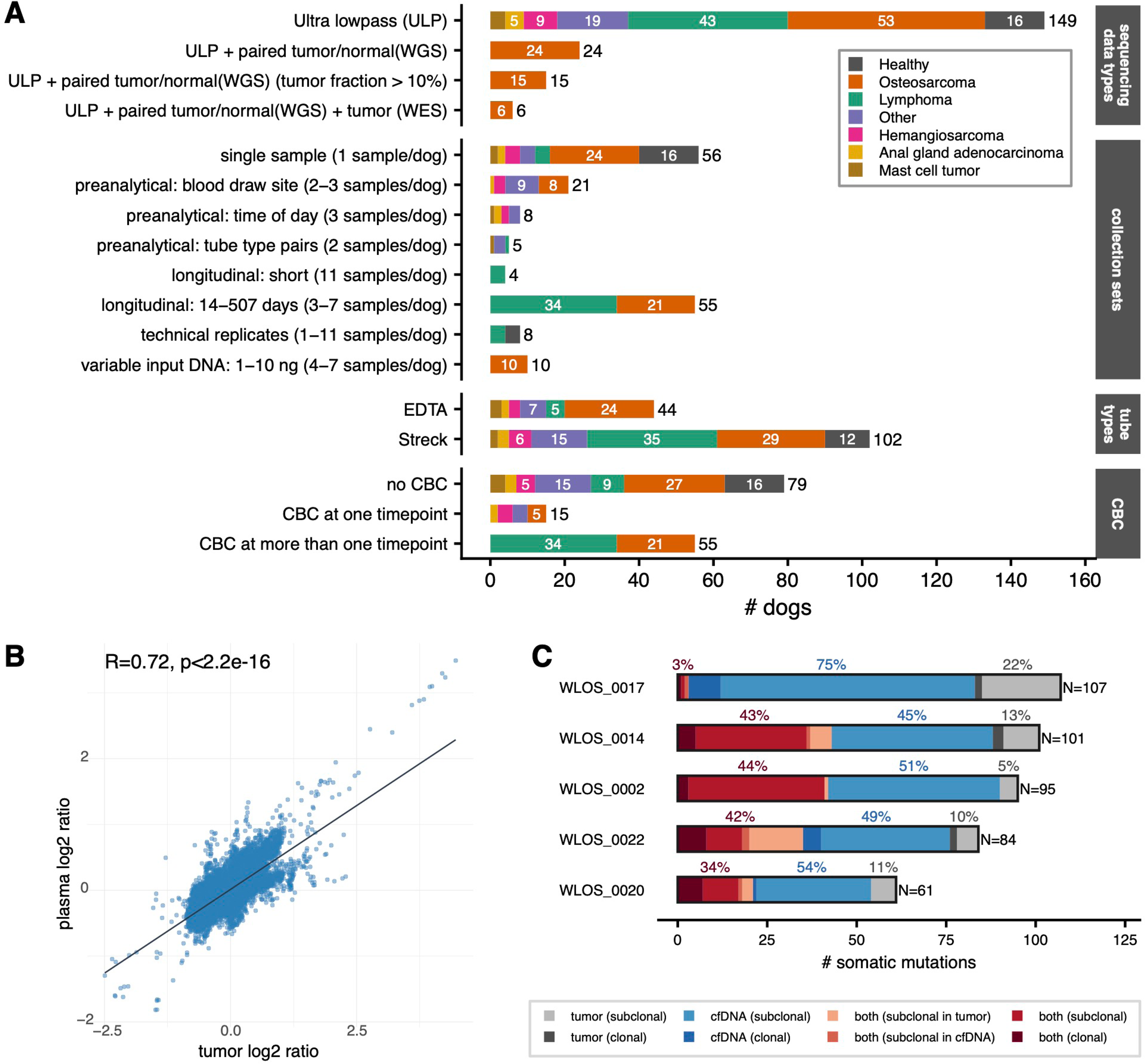
Overview of samples and tumor-cfDNA concordance. **(A)** Number of dogs broken down by number of samples and data types collected. **(B)** Spearman correlation of log2 read depth ratios between tumor WGS and cfDNA ULPWGS data in 14 dogs with cfDNA tumor fraction ≥ 0.1. **(C)** Proportion of somatic mutations identified in tumor, cfDNA, and shared between both in samples from five dogs.

The majority of liquid biopsy samples were collected in Streck cell-free DNA blood collection tubes (10 mL of blood) (Table S1). A smaller set of liquid biopsy samples were collected in EDTA tubes either prospectively or banked (Table S1). For all samples, plasma was separated from blood cells (plasma volume ranging from approximately 0.5 mL to 5 mL), and cfDNA isolated from the plasma. We then performed ultra-low-pass whole-genome sequencing (ULPWGS) of cfDNA at a target sequencing depth of 0.1x (Table SULPMETRICS).

We calculated three metrics from cfDNA samples and the resulting ultra-low-pass sequencing data: plasma cfDNA concentration (nanograms cfDNA per mL of plasma), tumor fraction, and genome-wide fragment size ratio. Tumor fraction is the fraction of the cfDNA derived from the tumor and was measured using ichorCNA adapted for use in dogs^10^. Fragment size ratio is the ratio of small (100 - 150 bp) to large (151 - 220 bp) DNA fragments as defined by DELFI Diagnostics^50^. A higher fragment size ratio indicates a greater proportion of short fragments, potentially reflecting a higher tumor fraction or increased apoptosis. Analyses reported below include only the first sample from each dog in our cohort unless multiple samples were part of the experimental design (*e.g.* blood draw site, time of day, longitudinal analyses). The first time point was dictated by clinical trial design, correlating with a pretreatment screening visit in a clinical trial unless otherwise stated (Table S1).

### Liquid biopsy metrics in dogs match human studies

Liquid biopsies in dogs and humans performed comparably on all three metrics (plasma cfDNA concentration, tumor fraction, and fragment size ratio). To enable comparisons of dog to human, we compiled human data from previously published studies (Table SHUMAN)^10,51^. Species had no significant effect on tumor fraction, and a small effect on cfDNA concentrations (ANOVA ges=0.008; p=0.014) and fragment size ratio (ges=0.1; p=1x10^-7^) (Fig SMETRICS A-D). In comparison, cancer type had a significant effect on all three: cfDNA concentration (ges=0.25; p=6.9x10^-36^), tumor fraction (ges=0.21; p=9.88x10^-22^) and fragment size ratio (ges=0.21; p=7.46x10^-7^). For example, while the average tumor fraction was lower in humans than in dogs (average of 0.14 ±0.183 in humans vs 0.184 ±0.194 in dogs; p_t-test_=9x10^-9^), this difference was reversed and nearly eliminated when we removed the dog lymphoma samples (average tumor fraction = 0.14 ±0.183 in dogs; p=0.05) (Fig SMETRICS E). While the effect of species on fragment size remains when limiting to canine EDTA samples only, some of the difference may be attributable to differences in sample handling and storage (including time to plasma separation), DNA extraction protocols, and library construction techniques^51^.

Technical variation in the amount, concentration and fragment sizes of the tumor DNA detected in the cfDNA samples was extremely low, with all three measures significantly and strongly correlated in 47 pairs of technical replicates (Fig SREPLICATES F-H). Consistent with benchmarking in human samples^52^, tumor fraction estimates were also strongly correlated (R=0.99, p=8x10^-17^) between libraries made using higher and lower DNA inputs from 32 cfDNA samples from dogs with osteosarcoma. Fragment size ratios were less strongly correlated across input replicates (R=0.792, p=1x10^-4^), suggesting that this metric may be more susceptible to input DNA amount (Fig SINPUT). As expected, the tumor fraction was positively correlated with the plasma cfDNA concentration in liquid biopsies from dogs with cancer (R=0.58; p=2.5x10^-11^; N=112) but not in healthy dogs (R=0.14; p=0.65; N=14). cfDNA concentration was negatively correlated with fragment size ratio, especially in dogs without cancer (R=−0.625, p=0.0072) (Fig SCOMPARE).

### cfDNA samples capture somatic mutations missed in sequencing of matched tumor tissue

Liquid biopsies in dogs capture somatic changes observed in tumor tissue collected from the same dog at the same time, consistent with findings in humans^53^. For 24 dogs with osteosarcoma^54^, we identified large somatic copy number changes in the tumor (60x WGS) and the cfDNA (ULPWGS) compared to normal muscle tissue (30x WGS) using ichorCNA (Table_STF_SAMPLETYPE). Copy number changes were significantly correlated between tumor and matched cfDNA samples (Spearman’s rho = 0.63; p<2.2x10^-16^, N=24). The correlation between sample pairs with cfDNA tumor fractions ≥ 0.1 was even stronger (rho=0.72; p<2.2x10^-16^; N=14) and nearly identical to that observed in humans (0.763 in 22 sample pairs of metastatic breast cancer^10^) (Fig 2B, Fig SSPEAR, Table_SREADDEPTH).

To investigate whether simple somatic mutations (SNVs and indels) are captured by liquid biopsies, we developed a custom whole exome sequencing (WES) panel^55^. Our design targets 37.6 Mb and includes 40.5 Mb of probes covering 20,257 genes as well as non-coding regulatory regions implicated in human cancers. The probes are available as a product through Twist Bioscience as a resource for other comparative canine genomics studies. For six of the 24 osteosarcoma samples, we performed WES to an average target coverage of 131x (Table SWESMETRICS). We called somatic mutations using Mutect2^56^, Strelka2^57^, and Octopus^58^ and kept mutations identified by two or more callers for further analysis. One sample was excluded from further analysis because there was evidence of cross-sample contamination.

We assessed the concordance of clonal and subclonal simple somatic mutations between tumor and paired cfDNA samples (Table SCONCORDANCE). Clonal mutations were defined as having an allele fraction greater than or equal to 0.9 times the tumor fraction as estimated by ichorCNA. Concordance between tumor and cfDNA was high for four sample pairs, with an average of 88% of clonal mutations found in the tumor also seen in the cfDNA (86%, 92%, 100%, and 100%) and an average of 86% of clonal cfDNA mutations also observed in the tumor (66.7%, 88.9%%, 100%, and 100%). Concordance of subclonal mutations was lower, averaging 79% for tumor mutations found in cfDNA, and 39% of cfDNA mutations found in the tumor. One sample (WLOS_0017) had much lower concordance, with just 33% of clonal and 8.3% subclonal tumor mutations seen in the cfDNA, and 18.2% of clonal and 1.4% of subclonal cfDNA mutations seen in the tumor. The cause of the low concordance in this sample is unknown. Tumor fractions (tumor: 0.6, cfDNA: 0.28) were within the range observed in the other samples, and we confirmed that the tumor and cfDNA were from the same individual using Bam-Matcher^59^ (Table SCONMATCH, Fig_SSWAP). In all five cfDNA/tumor pairs, we detected a larger number of somatic mutations in cfDNA WES than in tumor (74 vs. 34 variants on average, paired one-tailed t-test p=1.92x10^-4^) suggesting that cfDNA could be capturing a more complete representation of tumor simple somatic mutations than single-tumor biopsies (Fig_2C). This aligns with previous work in humans^60^. Consistent with this, the cfDNA also tended to have lower allelic fractions (paired t-test p=0.013) and higher MATH (Mutant-Allele Tumor Heterogeneity)^61^ scores (paired t-test p=0.05) compared to the tumor (Fig SCONCORDF,G, Table SCONMATH).

### Liquid biopsy metrics are affected by clinical variables and preanalytic factors

Dogs with cancer had significantly elevated values on all three cfDNA metrics compared to healthy controls (Fig 3 A-C), consistent with human studies^10,51^. About half of the variation in liquid biopsy metrics was explained by differences among individual dogs (Fig 3 D), which could reflect variation in clinical features, other lifestyle factors, or inherited genetics. The effect of the individual identifier was between 0.49 and 0.58 for all three performance metrics. None of the patient characteristics, including breed, sex (with spay/neuter status), weight, or age, explained the effect, consistent with results from human studies that found a weak or mixed effect of age, sex, and weight^62^.

**Figure 3.**
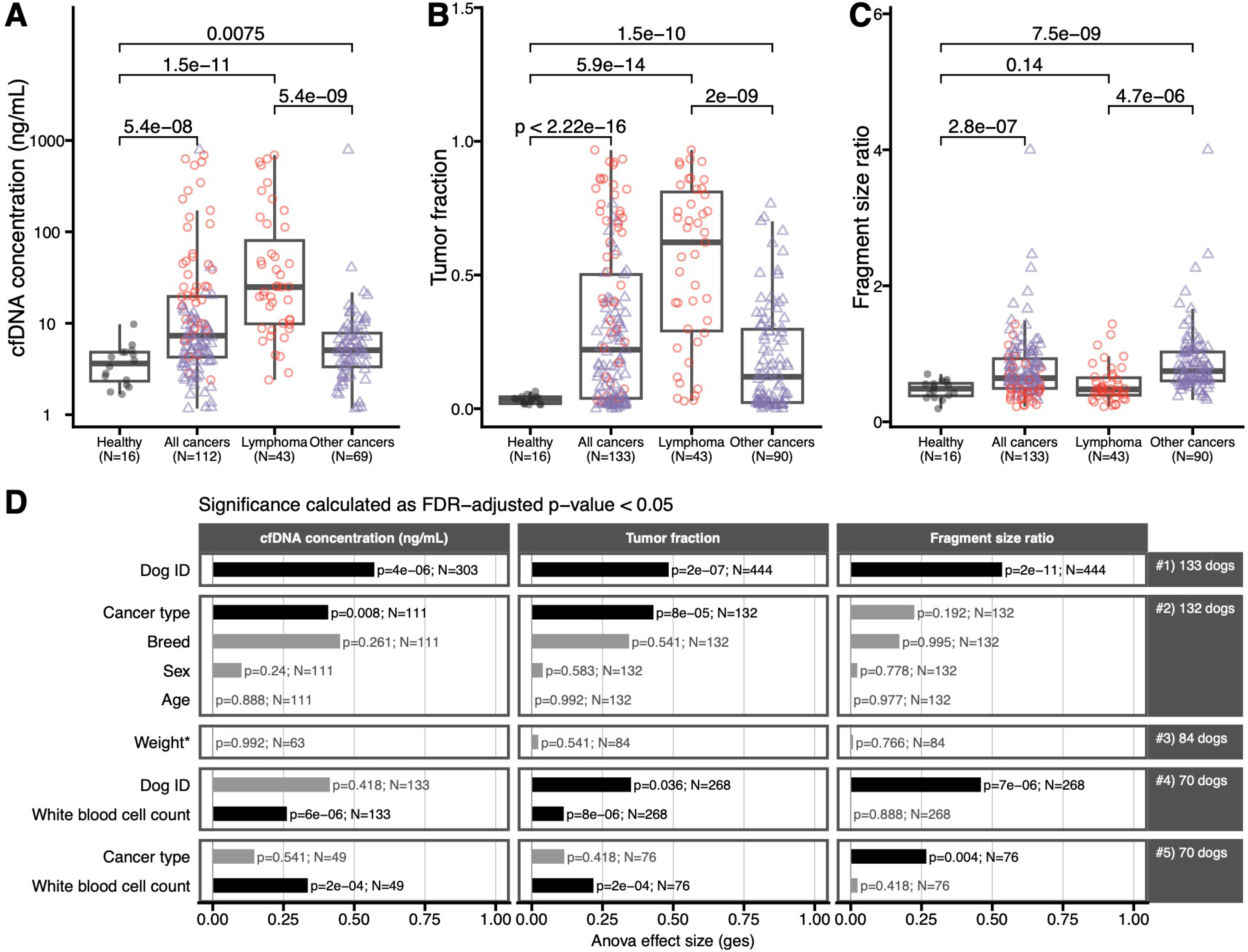
Patient factors affecting cfDNA metrics. **(A)** cfDNA concentration in healthy dogs, all cancers, lymphoma, and other cancers. **(B)** cfDNA tumor fraction in healthy dogs, all cancers, lymphoma, and other cancers. **(C)** cfDNA fragment size ratios in healthy dogs, all cancers, lymphoma, and other cancers. All metrics are significantly different between healthy dogs and those with any cancer, and between lymphoma and other cancers, with lymphomas tending to have higher cfDNA concentrations and tumor fractions but lower fragment size ratios. (D) ANOVA results showing the effect of various patient factors on cfDNA metrics. Dog ID had the largest effect on all metrics. Cancer type had a significant effect on cfDNA concentration and tumor fraction. White blood cell count had a significant effect on metrics both independently and when controlling for cancer type.

### In dogs with cancer, histologic type has a large effect on liquid biopsy metrics

Apart from presence or absence of cancer, the feature with the strongest effect on cfDNA metrics was the type of cancer (Fig 3 D). Dogs with lymphoma had higher cfDNA concentrations and higher tumor fractions than either healthy controls or dogs with other types of cancer (Fig 3 B-D). This could potentially be due to differences in stage between the cancer types. The majority of dogs with lymphoma were enrolled as part of a clinical trial with stage III or IV disease (meaning that more systemic signs of disease were present). In contrast, the majority of dogs with osteosarcoma had a primary tumor without detectable metastasis at enrollment (more localized disease). Sample collection from dogs with other cancer types was not standardized to a specific treatment time point. Because stage is largely confounded with cancer type in this study, we cannot discern which difference is causal. Other characteristics of individual dogs did not significantly impact cfDNA metrics. The effect size of dog breed was large, but did not reach significance in the statistical model overall. Because few breeds were represented by more than one dog in our data set (Fig SBREEDMETRICSA), we could not distinguish a potential effect of these breeds from other characteristics that differ between individuals. Comparing the two most highly represented breeds (Labrador retriever and golden retriever) and all other dogs revealed no significant differences except for a significantly lower fragment size ratio in Labrador retrievers than other breeds (p=0.0052) (Fig SBREEDMETRICSB-G).

White blood cells are a primary source of cfDNA, even in healthy individuals, so it is possible that differences in metrics for lymphomas as compared to other cancer types are driven by the elevated (neoplastic) white blood cell counts characteristic of this cancer type^63,64^. Complete blood count (CBC) data was available from 70 dogs with cancer (34 with lymphoma, 26 with osteosarcoma, 10 other cancers), in some cases from multiple time points, for a total of 270 samples (Table SCBC). Analyzing the first sample (usually pre-treatment at enrollment in a clinical trial) per dog only, the effect of lymphoma on concentration of cfDNA and tumor fraction was largely explained by the higher number of white blood cells detected in lymphoma samples (Fig 3D). In this cohort, white blood cell count was positively correlated with cfDNA concentration in all samples, although this was only significant in dogs with lymphoma (R=0.604, p<2x10^−4^). Suggestive of the involvement of neoplastic lymphocytes, white blood cell count was positively correlated with tumor fraction only in lymphomas (R = 0.615, p=1x10^−4^), and negatively correlated with fragment size ratio (R=−0.40, p = 0.02 (Fig 4A-C). These findings suggest that in dogs with lymphoma DNA released from the circulating neoplastic white blood cells either *in vivo* or during the sample collection process may increase both the cfDNA concentration and the fraction of cfDNA that is tumor derived, and that the tumor DNA from these cells is less fragmented than might be expected if it had traveled from a non-blood-cell source.

**Figure 4.**
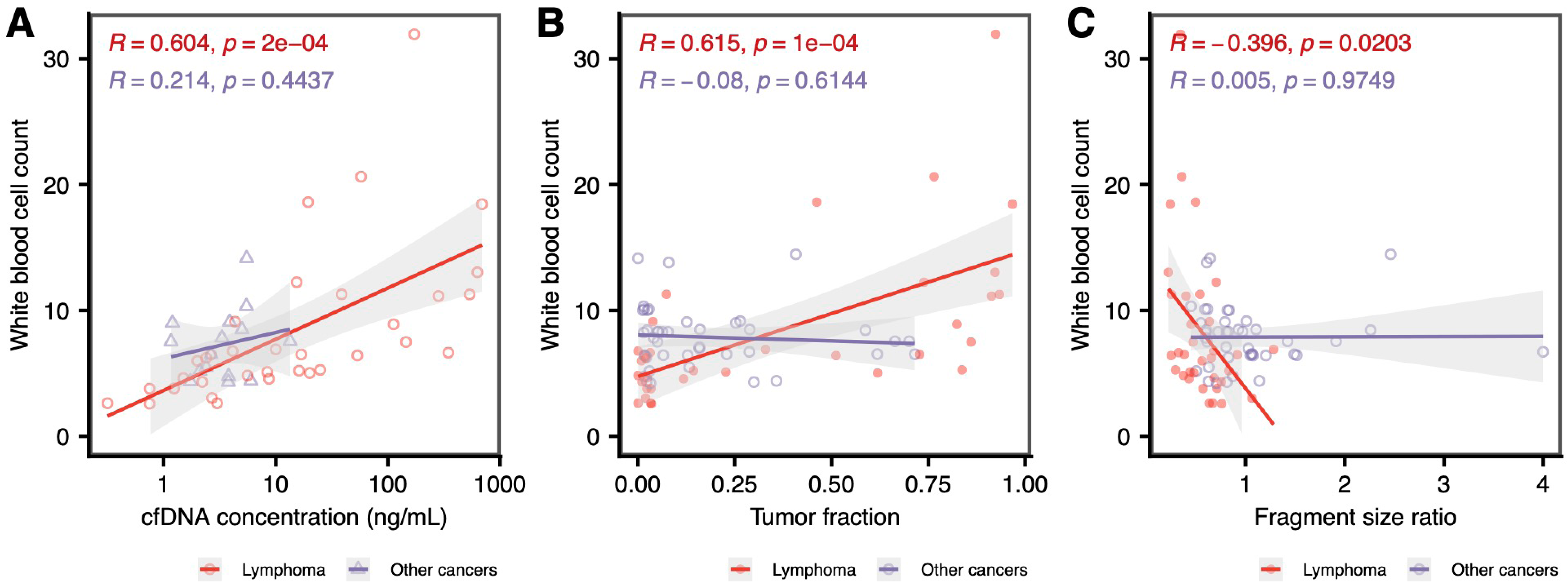
Correlation of white blood cell count to cfDNA metrics. Considering the first sample only from each dog with complete blood count data, white blood cell count was significantly positively associated with **(A)** cfDNA concentration and **(B)** tumor fraction, and significantly negatively associated with **(C)** fragment size ratio in lymphoma but not other cancers.

### Blood collection tube type affected sample quality

Whether a sample was collected in a conventional EDTA tube or a specialized blood collection tube has a small effect on the amount and quality of cell-free DNA collected. Streck tubes are designed for liquid biopsy applications and contain specialized preservatives to prevent cell lysis^14^. For five dogs, each with a different type of cancer, we collected into both EDTA and Streck tubes at the same visit. There was no effect of tube type on the detected tumor fraction (mean_EDTA_= 0.16, mean_Streck_=0.17, paired p_t-test_=0.22), but fragment length ratios were modestly higher in the Streck tube samples (smaller fragments) (mean_EDTA_= 0.80, mean_Streck_=0.88, paired p_t-test_=0.0072), potentially consistent with decreased white cell lysis or inhibition of DNase activity in the Streck samples^23^ Fig STUBES). We were not able to assess the effect of tube type on cfDNA concentration because we didn’t have total plasma quantities for these five samples, so we expanded this comparison to consider all samples where the blood collection tube was known. After excluding the dogs with lymphoma, for which samples were predominantly collected into Streck tubes (N=34) rather than EDTA tubes (N=4), we saw an effect of tube type on the concentration of cfDNA. Streck tubes yielded significantly lower concentrations (p_t-test_=6x10^-8^) potentially indicative of less white cell lysis. Most samples collected in EDTA tubes were processed within 30 minutes, however, in some cases EDTA samples from our pilot were fractionated up to 24 hours post collection Fig STUBES). The observation that dogs with lymphoma have higher cfDNA concentrations and tumor fractions was not attributable to tube type.

### Blood draws from a central and more tumor-proximal vein yielded more tumor-derived cfDNA

Given the short half-life of cfDNA in the circulation, we asked whether the proximity of the venipuncture site to the tumor or other characteristics of the central or peripheral vasculature might affect the amount of tumor-derived cfDNA sampled (and therefore the diagnostic quality of the liquid biopsy). This is difficult to assess in human patients, where a single sample is generally drawn at any given time point, and the venipuncture site is standardized (typically drawn from the median cubital or cephalic veins^25^). Despite this being a potentially simple procedure to optimize, only a few case reports in the human literature exist for adjusting sampling site as a way to increase tumor-derived cfDNA yield^65^, and none of these reports assessed tumor fraction.

To assess whether the vein used for liquid biopsy sampling affects liquid biopsy metrics, we collected blood samples from three different veins (jugular, cephalic, and saphenous veins) in the same dog at a single time point in a cohort of 10 dogs with various cancer types, and collected blood samples from two different veins (jugular and either cephalic or saphenous) at a single time point in a cohort of nine dogs (Table SSITE). All blood draws were performed using the same phlebotomy procedures, via percutaneous venipuncture with a sterile intravenous needle (20-22 gauge). We mapped the approximate tumor and blood draw site locations onto a standardized canine anatomy diagram and measured the distance between the tumor and blood draw sites. We considered both the “simple” straight-line distance and the “circulation” distance, which traces the path of the blood flow through the circulatory system from the tumor to the blood draw site.

Both jugular venipuncture site (p=1.9x10^-6^) and shorter circulation distances (p=1.4x10^-3^) were significantly associated with higher tumor fraction, even after controlling for inter-dog variability (Fig 5A, Fig SDRAW A). Because the jugular vein was always closer to the tumor than the saphenous vein (p_TukeyHSD_=0) or the cephalic vein (p_TukeyHSD_=6.3x10^-6^) (Fig 5B), the effect of sampling from a central vein could not be assessed separately from circulation distance. The average difference in tumor fraction between the closest site and the furthest site in samples with at least one nonzero tumor fraction was -0.043 (+/-0.039), and we recovered an average of 0.3-fold (+/-0.3) lower tumor fraction from the furthest site than the closest. Jugular vein site was also associated with higher fragment size ratios (overall smaller fragments), p=2.3x10^-4^ (Fig 5C), while circulation distance was not significantly associated with fragment size ratios (p=0.17) (Fig SDRAW B). The concentration of cfDNA was not significantly associated with either jugular site (p=0.2) (Fig 5D, Fig SDRAW C) or circulatory distance (p=0.6) (Fig SDRAWB), suggesting that the central, closer site was enriched for tumor DNA. Simple distances (which did not take the path of blood circulation into account) were not significantly associated with liquid biopsy metrics (p=1 for all metrics).

**Figure 5.**
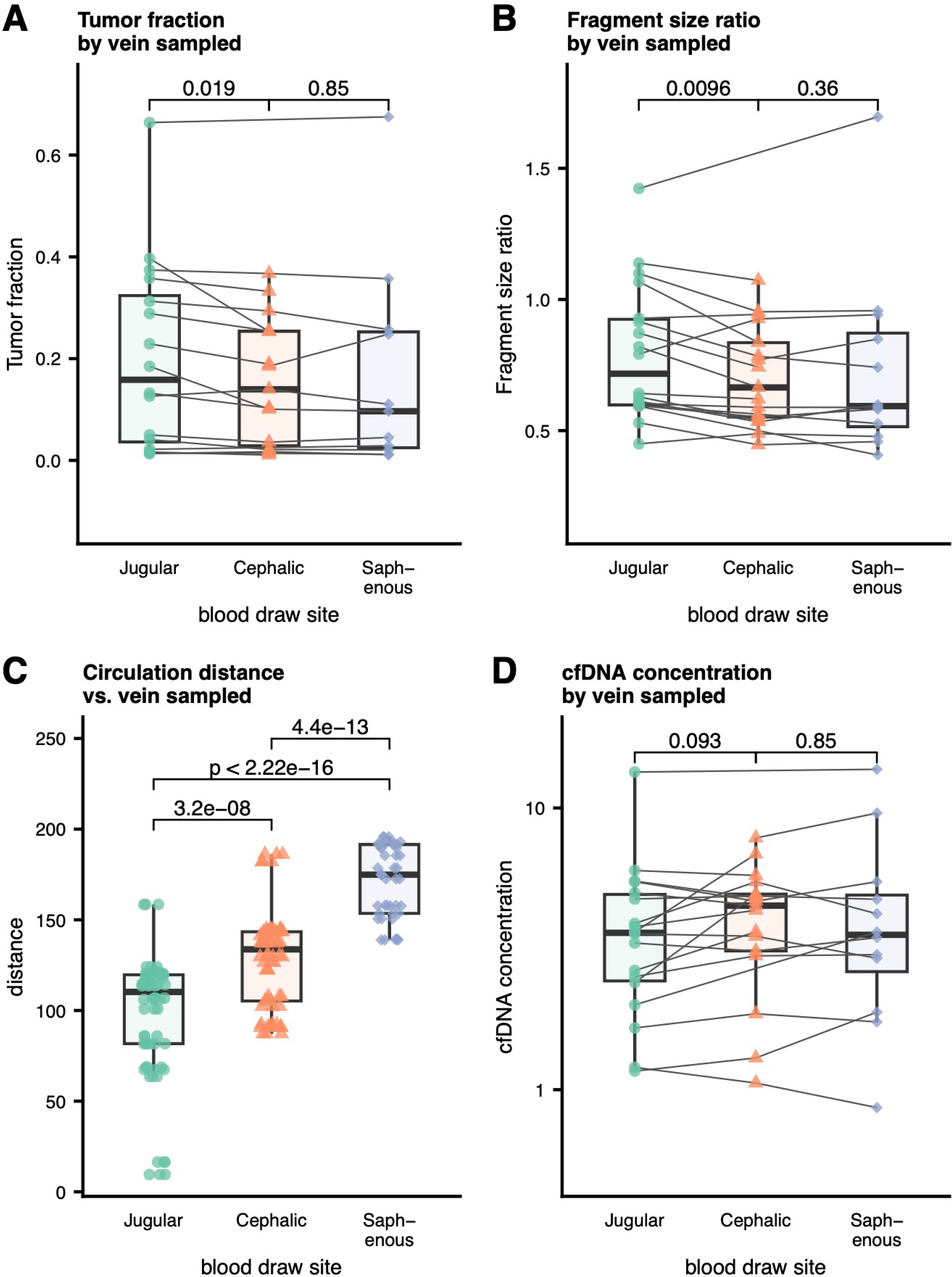
Effect of blood draw site on cfDNA metrics. **(A)** Jugular vein blood draw site was associated with higher tumor fractions. **(B)** The jugular vein was always the closest site to the tumor by circulation distance. **(C)** Jugular vein site was significantly associated with fragment size ratio. **(D)** cfDNA concentration was not significantly associated with blood draw site.

### Time of day had no significant effect on cfDNA metrics

Studies in humans have not conclusively determined whether the time of day of sampling has an effect on cfDNA yields^16,17,66,67^. However, circadian rhythm and sample timing has been shown to be important in blood cell^16,68,69^ and circulating tumor cell counts^70^. We examined the effect of the time of blood sampling in a cohort of 10 dogs with various cancer types. Among three samples taken at intervals of 2 to 3.5 hours (Table STIME), we found no significant effect of time of day on either cfDNA concentration (p=0.9), tumor fraction (p = 0.9), or fragment size ratio (p=0.5) (Fig STIME A-C). We also saw no difference between morning and afternoon blood draws on any metric (Fig STIME D-E). It is possible, however, that longer sampling intervals (*e.g.,* overnight) would show a more significant effect.

### Longitudinal cfDNA metrics correlated with disease status

Detection of molecular relapse is an important clinical application of liquid biopsy and is under active development^71,72^. It is estimated that a tumor must reach a size of approx 1.5 cm in diameter (approx 1.77 mL volume^73^) before it is detectable by clinical imaging, thus, using liquid biopsy to identify molecular relapse before the tumor is visible on imaging may improve patient outcomes. Such studies take years in human patient cohorts, but can be performed much more quickly in canine patients, who have an accelerated clinical course and shorter lifespans. In order to evaluate the comparative utility of the dog model to help optimize longitudinal patient monitoring, we assessed cfDNA dynamics and dog clinical status over short (hours to days) and long (weeks to months) timescales. We collected multiple cfDNA samples over the course of treatment from dogs with lymphoma or osteosarcoma enrolled in clinical trials at the Tufts Foster Hospital for Small Animals, noting, at each time point, disease status (whether gross disease was present and whether there was a response to therapy) based on patient clinical status and RECIST criteria^74^, which standardize the designation of response based on changes in tumor size. Samples collected longitudinally at appointments where treatment was administered were always taken pre-treatment. We did not collect paired tumor biopsies, although in some cases samples were collected prior to surgical biopsy or lymph node aspiration for clinical use.

### cfDNA metrics change within hours of L-asparaginase treatment

Little is known from either mouse models^75^ or human patients^76^ about the dynamics of cfDNA over short time frames after treatment. Time scales for blood collection are often on weekly to monthly or even quarterly intervals whereas the half-life of cfDNA is on the order of minutes to hours^77^. Identifying the post-treatment interval in which liquid biopsy samples have the highest tumor content, or in which cfDNA metrics are most predictive of response could help to maximize tumor-derived cfDNA yield and provide an early assessment of treatment efficacy. To examine cfDNA dynamics over short (6 hours to 1 week) time periods, we recruited four dogs with lymphoma (three with multicentric diffuse large B-cell lymphoma (DLBCL) and one with epitheliotropic lymphoma). For each dog, we took blood samples 6-12 hours apart during the 24 hours prior to treatment with one dose of L-asparaginase, then again during the 24 hours following treatment, and, finally, during the 24 hour period one week later. We assessed the relationship between cfDNA metrics from the three multicentric samples and the sum of the five lymph node sizes measured for RECIST^74^ staging each day (in the three dogs with DLBCL), as well as the clinical assessment of the attending veterinarian (Table SARM).

Relative changes in all three metrics qualitatively followed the clinical responses of the dogs (Fig. 6, Fig SSHORT). Fragment size ratios increased by 12 hours post-treatment in all dogs (Fig 6D), consistent with an increase in apoptosis and tumor DNA release into the circulation immediately post-treatment. For three of the dogs, fragment size ratios then decline, while in the dog who had a good clinical response overall, fragment size ratio remained elevated at 48 hours post-treatment (Fig 6D). These findings suggest that very early post-treatment changes and kinetics may be predictive of response in a way that a single time point one week post-treatment would not be, although a larger cohort will be needed to further assess short-term cfDNA kinetics post-treatment.

**Figure 6.**
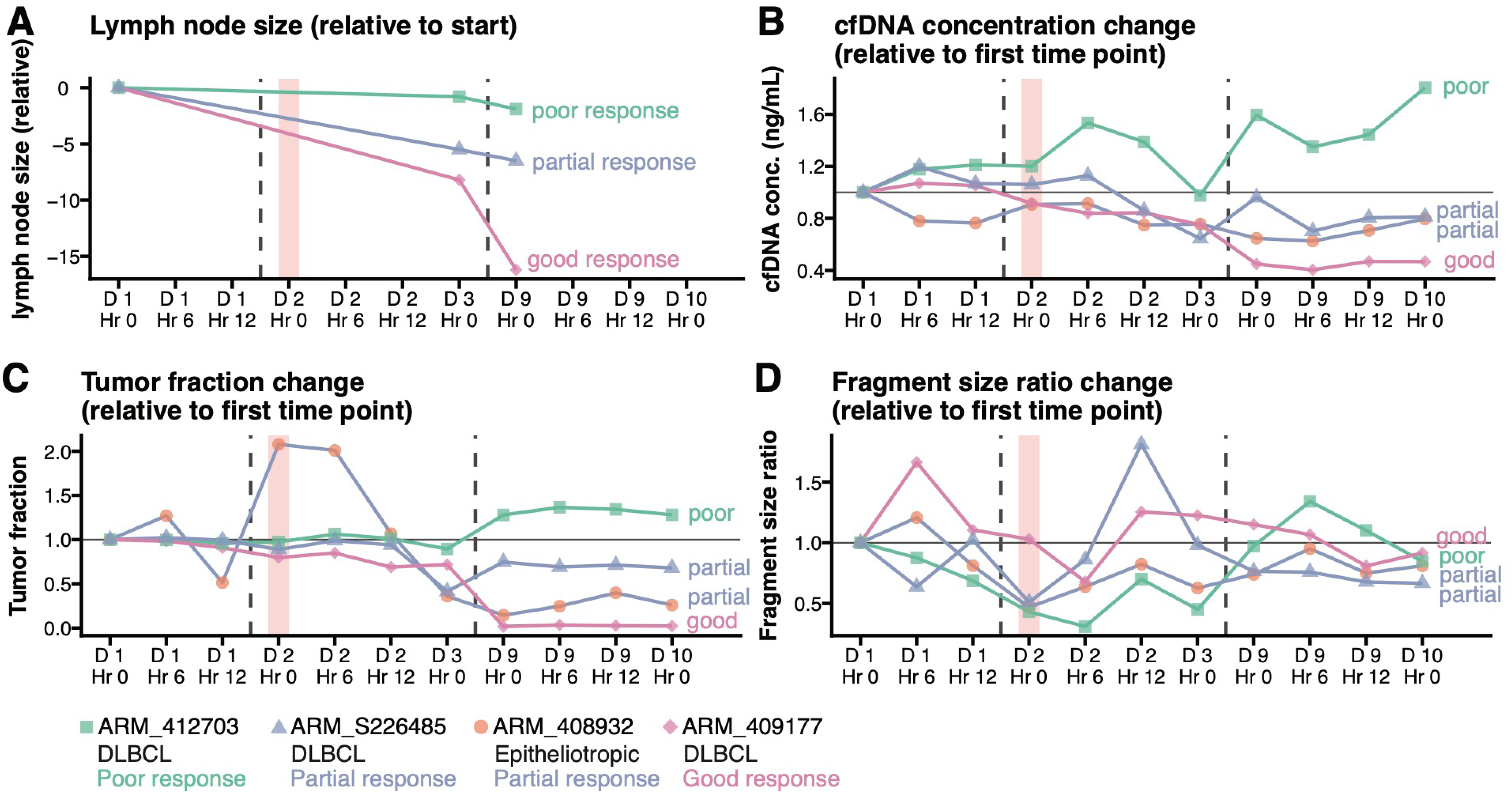
Acute response monitoring. Short-term cfDNA kinetics in four dogs with lymphoma with samples taken over 24 hours pre- and post-treatment with L-asparaginase (treatment time point indicated as pink bar) and 24 hours one week later (dotted lines indicate break between days. **(A)** Lymph node size on days 1, 3, and 9 in three dogs with DLBCL. **(B)** Relative change in cfDNA concentration over treatment. **(C)** Relative change in tumor fraction over treatment. **(D)** Relative change in fragment size ratio over treatment. All dogs had an increase in fragment size ratio by 12 hours post-treatment.

### Changes in cfDNA metrics are detected prior to clinical relapse in lymphoma and osteosarcoma

The clinical utility of using liquid biopsy metrics, including tumor fraction and cfDNA concentration, for monitoring molecular response to assess the effectiveness of therapy and as predictors for outcomes such as progression-free survival in human oncology is a promising and active area of investigation^78,79^. We therefore asked how well liquid biopsy values compare with clinical metrics across a full course of cancer treatment, which can be completed in much shorter time frames than human studies — approximately one year for canine lymphoma and osteosarcoma patients.

Longitudinal data was collected from a total of 34 dogs with DLBCL and 21 dogs with appendicular osteosarcoma undergoing treatment as part of clinical trials at Tufts Cummings School of Veterinary Medicine (Table S1). Samples from 3 - 5 clinical visits were collected from the dogs with lymphoma, including screening/day 0, day 7 or 14, day 21 or 28, the visit prior to clinical relapse (defined retrospectively), and clinical relapse. A subset of six dogs had not yet relapsed, so the sample from the latest visit was analyzed. For dogs with osteosarcoma, samples from 3 - 8 time points were collected every two weeks for the first month (including screening visit and amputation at week 2), then monthly thereafter until relapse. Again, for a subset of 6 dogs that had yet to relapse, samples up to the latest time point collected were included.

We assessed pre- and post-amputation time points from the 21 dogs with osteosarcoma to benchmark the ability of tumor fraction to identify a clinically well-defined decrease in tumor burden. The median change in tumor fraction was 0.135 +/-0.154, with 91% of dogs showing a decrease in tumor fraction (SAMP). We noted similar trends in fragment size ratios in 71.4% of dogs. Because all dogs underwent amputation to remove the primary tumor, we were unable to compare these trends between dogs that had the intervention and those that did not.

cfDNA metrics were significantly associated with disease status across all time points in both cancer types. In lymphoma all three metrics were significantly associated (logistic regression, log10 cfDNA concentration p=0.00209; tumor fraction p=0.02575, fragment size ratio p=0.04061). In osteosarcoma tumor fraction was significantly associated with disease status (p=0.00215) (Fig SLONGDOG_LSA_CONC, Fig SLONGDOG_LSA_TF, Fig SLONGDOG_LSA_FRAG, Fig SLONGDOG_OS_TF, Fig SLONGDOG_OS_FRAG).

We used paired t-tests to compare initial presentation (“disease present”) to intermediate time points with no clinical evidence of disease (“no evidence of disease”) and progressive disease (Fig 7, Fig SLONBOX). Dogs were considered to have “no evidence of disease” if they achieved a complete response and did not yet have progressive disease based on lymph node measurements (lymphoma) or if they were post-amputation and no metastases had been detected clinically (osteosarcoma). Full staging via diagnostic imaging was not performed. In some cases, elevated cfDNA metrics were identified at time points where dogs had no clinical evidence of disease, suggesting that minimal residual disease (MRD) was being detected at these time points. Between the “disease present” and “no evidence of disease” time points, where cfDNA metrics would be expected to decrease, cfDNA concentration significantly decreased in lymphomas (p=1.2x10^-6^) (Fig 7A), tumor fraction significantly decreased in lymphoma (p=2.4x10^-8^) and osteosarcoma (p=1.4x10^-5^), and fragment size ratio significantly decreased (p=0.0055) in osteosarcoma. Between the “no evidence of disease” and “progressive disease” time points, where cfDNA metrics would be expected to increase, we found that cfDNA concentration (p=0.0018) and tumor fraction (p=0.00025) significantly increased in lymphoma (Fig 7B), while fragment size ratio significantly increased in osteosarcoma (p=0.02) (Fig 7C). These increased cfDNA metrics suggest that MRD was increasing with disease burden and molecular evidence of relapse was detectable at the clinical relapse time point.

**Figure 7.**
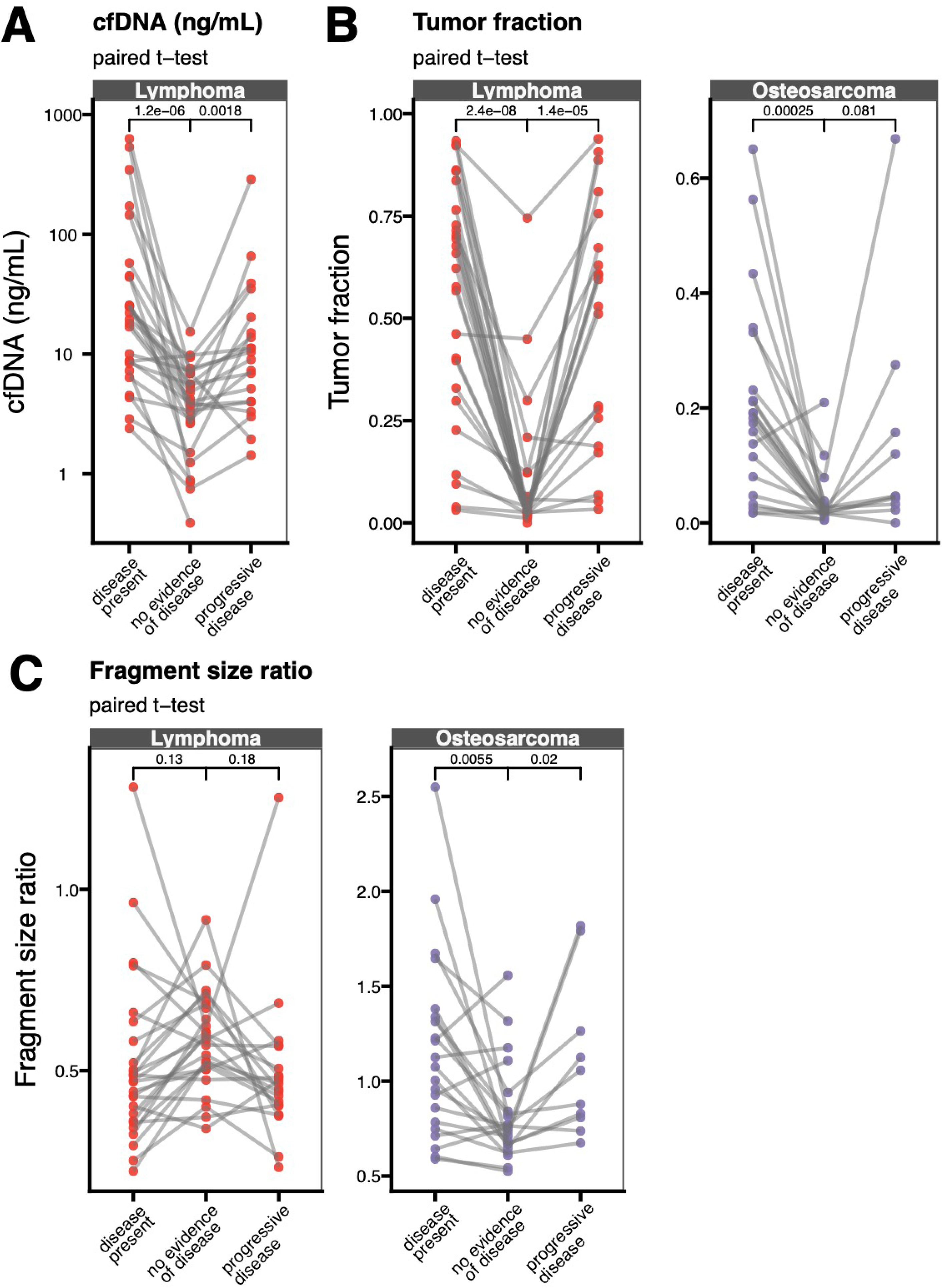
Longitudinal monitoring. Multiple samples from dogs with lymphoma or osteosarcoma over the course of treatment. Plots compare initial pre-treatment of pre-amputation time points (“disease present”), remission or post-amputation time points (“no evidence of disease”), and “progressive disease” time points. **(A)** cfDNA concentration was associated with disease status in dogs with lymphoma. **(B)** Tumor fraction was significantly associated with disease status in lymphomas, and between “disease present” and “no evidence of disease” in osteosarcomas. **(C)** Fragment size ratio was significantly associated with disease status in osteosarcomas but not lymphomas.

To investigate whether evidence of molecular relapse prior to clinical relapse could be identified, we assessed trends in cfDNA metrics from screening to relapse. We identified 17 dogs (9 with lymphoma and 8 with osteosarcoma) with four or more longitudinal samples, including one time point with no evidence of disease and the clinical relapse time point (with at least one intervening time point) (Table STREND). We found that there were patterns of increasing tumor fraction with increases continuing at each time point from initial increase until clinical relapse. This pattern of increasing tumor fraction was seen in 8/9 (89%) of dogs with lymphoma and 5/8 (63%) of dogs with osteosarcoma. These trends started an average of 41 days prior to clinical relapse in lymphoma and 30 days prior in the osteosarcomas, although it should be noted that the sampling interval was dictated by the frequency of clinical trial visits. We identified similar trends in cfDNA concentration in lymphomas, which increased in 2 dogs an average of 30 days earlier than clinical relapse, and in fragment size ratios, which increased in 2 dogs with osteosarcoma an average of 30 days before clinical relapse, suggesting that these metrics may also be useful for patient monitoring (Table STREND).

Using Cox proportional hazards models, we investigated whether cfDNA metrics were predictive of outcome. In lymphomas, the log_10_ cfDNA concentration was associated with progression-free survival (PFS) (p=0.015) at the baseline time point (Fig SFOREST A), while at day 7, when all dogs had achieved either a complete or partial response, tumor fraction was significantly associated with PFS when controlling for disease status (p=0.013) (Fig SFOREST B). Baseline and post-amputation time points were not significantly associated with outcome in osteosarcoma. Our results lend weight to previous canine studies that tumor-derived cfDNA levels can be predictive of outcome^34,35,44^, identify residual disease after curative-intent surgery^43,46^, and predict relapse when patients were monitored longitudinally with serial liquid biopsies^39,45^.

## DISCUSSION

In this study, we demonstrated that spontaneous canine cancer is a powerful comparative model for optimizing liquid biopsy techniques. To validate the canine model, we confirmed that several key metrics for dog cancers— including tumor-cfDNA concordance, cfDNA plasma concentration, tumor fraction, and fragment size ratios—closely match previous reports from human patients. Leveraging the higher incidence of cancer types such as lymphoma and osteosarcoma in dogs, their rapid clinical course, and flexibility in veterinary (as compared to human) phlebotomy and treatment protocols, we identified factors affecting cfDNA metrics that have not been reported previously or that are not yet commonly considered in liquid biopsy optimization. These include differences between lymphoma and nonhematopoietic cancers and the impact of central venous sampling and circulatory distance on tumor fraction.

While numerous studies have demonstrated the feasibility of liquid biopsy in pet dogs with cancer^34–40,42–49,80^, ours is the first to specifically evaluate canine liquid biopsy performance in the context of comparable human studies. We found that the ranges of cfDNA metrics in dogs with and without cancer were similar to metrics in humans with and without cancer, that concordance of somatic mutations between tumor and cfDNA samples was very similar to that reported in humans^10^, and that more mutations are identified in cfDNA than in tumor tissue, as has been reported in humans^60^. Taken together, our findings indicate that liquid biopsy performance is similar in dogs and humans, verifying that spontaneous cancers in dogs are an excellent model system for exploring novel methodologies to advance liquid biopsy performance and accuracy.

We identified key differences between cfDNA metrics between lymphomas and non-hematopoietic cancers that point to cancer type as an important consideration in clinical interpretation of liquid biopsy findings.

Overall, lymphomas tended to have higher cfDNA concentrations and tumor fractions but lower fragment size ratios than non-hematopoietic cancer types. Total white blood cell count was associated with higher cfDNA concentration and tumor fraction but lower fragment size ratios in lymphomas, but not other cancer types. These findings suggest that while cfDNA from normal white blood cells dilutes cfDNA with normal germline DNA, in individuals with lymphoma, circulating neoplastic lymphocytes instead contribute tumor DNA. It has been reported that tumor cfDNA tends to be more highly fragmented than normal cfDNA^81^. Our findings suggest cfDNA contributed by white blood cells directly in the bloodstream is less fragmented than DNA from extravascular sources.

Our finding that the circulatory proximity of the blood draw site to the tumor is important for understanding cfDNA kinetics. Because the jugular vein was always the closest site in our study, further research will be necessary to identify whether tumor fraction decreases with distance of the blood draw site from the tumor, or whether other properties of central venous sampling (including aggregation of blood from large areas of the peripheral circulation, and potentially from unknown metastases) increase tumor cfDNA yield. There may also be cases in low-cfDNA cancers for selective venous sampling of more tumor-proximal vasculature - this approach has already been adopted in non-liquid biopsy approaches including localization of endocrine disease^82^, and it has been shown to be more sensitive for quantifying prostate specific antigen^83^ and capturing circulating tumor cells^65,84,85^. A recent study reported higher concentrations of cfDNA and more reliable identification of tumor mutations from blood samples more proximal to the tumor in three human patients^65^, suggesting that selective venous sampling could lead to improved tumor-derived cfDNA yield and assay performance in humans as well.

This is the first study to evaluate cfDNA at such dense time points pre- and post-treatment. We utilized this data to identify a pattern in fragments size ratios post-treatment that may indicate increased levels of treatment-induced apoptosis. In addition, the longitudinal studies across treatment of dogs with lymphoma and osteosarcoma are among the largest to date in single cancer types. Our findings confirm the utility of liquid biopsy for identifying molecular relapse prior to clinical detection of cancer, and benchmarks a lead time for when emergence of therapeutic resistance in osteosarcoma and lymphoma can be reliably ascertained.

In conclusion, our research illuminates the potential of cell-free DNA (cfDNA) as a transformative model for liquid biopsies for cancer detection and monitoring in human and veterinary medicine. In addition to identifying key factors for future optimization studies, this work positions spontaneous cancers in pet dogs to be more broadly employed for comparative studies designed to innovate, optimize and validate liquid biopsy methodologies prior to human translation.

## METHODS

### Ethical approval

Ethical approval for sample collection was obtained from the Institutional Animal Care and Use Committees (IACUCs) at Tufts Cummings School of Veterinary Medicine (TCSVM), The Ohio State University School of Veterinary Medicine, and the Broad Institute of MIT and Harvard. All samples were collected by veterinary professionals with owner consent.

### Blood sampling, plasma fractionation, and storage

All blood samples were collected by veterinary medical professionals using standard protocols. *The Ohio State University Biobank*. Blood samples were collected in EDTA tubes, and plasma was fractionated within 30 minutes and stored in 0.5 mL aliquots at -80°C. *TCSVM Comparative Pathology and Genomics Resource:* Blood was collected into a Streck cell-free DNA blood collection tube. Plasma was fractionated following the manufacturer’s protocol, and stored in approximately 1 mL aliquots at -80°C. *Broad Institute Genomics Platform*. Blood collected in Streck tubes was centrifuged at 1900 g for 10 minutes and plasma was transferred to a second tube before further centrifugation at 15000 g for 10 minutes. Plasma was stored at -80°C until cfDNA extraction. The amount of input plasma was quantified in order to calculate cfDNA concentration per mL plasma. Concentrations were estimated based on the number of approximately 1mL aliquots in the first blood draw site cohort.

### DNA extraction and QC

#### Cell-free DNA

*Blood Biopsy group.* Frozen plasma from TCSVM was thawed and centrifuged at 15000 g for 10 minutes. If a sample did not meet the preferred starting input (2.1mL), 1x PBS was added to bring the volume up. cfDNA was extracted using the QIAsymphony DSP Circulating DNA Kit according to the manufacturer’s instructions. The extracted cfDNA was quantified using the PicoGreen (Life Technologies) assay and then frozen at -20°C until ready for further processing. *Broad Genomics Platform.* If a sample did not meet the preferred starting input (6.3mL), 1x PBS was added to bring the volume up. cfDNA was extracted using the QIAsymphony DSP Circulating DNA Kit according to the manufacturer’s instructions. DNA was normalized to be within the range of 25-52.5 ng in 50 uL of TE buffer (10mM Tris HCl 1mM EDTA, pH 8.0) according to picogreen quantification. *TCSVM Comparative Pathology and Genomics Resource.* Frozen plasma (up to 4mL) was extracted using the Apostle MiniMax cfDNA extraction kit (Apostle, LLC) on a Biomek i7 Hybrid liquid handling system (Beckman Coulter) per manufacturer instructions. cfDNA was quantified using PicoGreen (Life Technologies) or Qubit high sensitivity 1x dsDNA assay, per manufacturer instructions. cfDNA was stored at -80C until library construction. *<u>Whole blood</u>: Broad Blood Biopsy group.* gDNA was extracted from ∼200 μL whole blood using the QIAsymphony DSP DNA mini kit. The extracted gDNA was sheared to 150 bp using a Covaris ultrasonicator, and quantified using the Qubit dsDNA HS kit (Invitrogen), then frozen at -20°C until ready for further processing.

#### ULPWGS library construction

Library preparation was performed using a commercially available kit provided by KAPA Biosystems (KAPA HyperPrep Kit with Library Amplification) and IDT’s duplex UMI adapters. Unique 8-base dual index sequences embedded within the p5 and p7 primers (purchased from IDT) were added during PCR. Enzymatic clean-ups were performed using Beckman Coultier AMPure XP beads. Library quantification was performed using the Invitrogen Quant-It broad range dsDNA quantification assay kit (Thermo Scientific) with a 1:200 PicoGreen dilution (*Broad Genomics platform)*, PicoGreen (Life Technologies) (*Broad Blood Biopsy group)*, or TapeStation (*TCSVM Comparative Pathology and Genomics Resource).* Libraries were sequenced on Illumina HiSeq 2500 rapid run (100 bp paired-end reads, *Broad Blood Biopsy group and Broad Genomics Platform*), HiSeq X (151 bp paired-end reads, *Broad Genomics Platform*), NovaSeq 6000 (151 bp paired-end reads, *Broad Genomics Platform*), or NextSeq 2000 (151 bp paired-end reads, *TCSVM Comparative Pathology and Genomics Resource*) with an overall mean coverage of 0.297x +/- 0.3105 (range: 0.0085x - 1.56x).

### Alignment and preprocessing of ULPWGS data

#### Broad Genomics Platform and Blood biopsy group

ULPWGS sequencing data was preprocessed following the GATK best practices. Sequencing data was aligned to the CanFam3.1 reference genome^86^ using bwa^87^, and duplicate reads were flagged using Picard Tools MarkDuplicates. *TCSVM Comparative Pathology and Genomics Resource.* ULPWGS sequencing data were demultiplexed using BCL convert (v2.4.0) and aligned to the CanFam4 reference genome^88^ using BWA Aligner (v1.1.4) on the Illumina basespace sequencing hub.

#### Fragment size ratios

Fragment size counts were calculated from bam files using the Deeptools2^89^ “bamPEFragmentSize” command. The ratio of the count of fragments between 100-150 bp (short) to the count of fragments between 151-220 bp (long) were calculated for each sample.

#### Evaluation of tumor fraction

The tumor fraction of cfDNA ultra low pass whole genome sequencing data was assessed using the ichorCNA^10^ tool, modified to produce plots of the canine genome, and to allow the manual definition of sample sex. HMMcopy^90^ was used to produce read depth counts in 1 Mb intervals across the genome. Reference files for GC and mappability correction were generated using the HMMcopy utils package (https://github.com/shahcompbio/hmmcopy_utils). Putative centromere locations were assigned as regions with 80% or more repetitive sequence, based on a genome scan for 4 centromeric repeats using a 5 kb window size (CanFam3.1: https://github.com/Chao912/Mischka/blob/master/CanFam3.1.centromere.bed; CanFam4: https://github.com/Chao912/Mischka/blob/master/GSD1.0_CanFam4.centromere.bed). The pseudoautosomal region was filtered out to prevent spurious copy number gains after chrX normalization in male dogs. As the first tumor fraction solution provided by ichorCNA is not always the best based on heuristics developed by the Broad Blood Biopsy team, we developed a set of rules to select the best solution in an unbiased way, prioritizing lower fraction SCNAs subclonal, lower copy number levels, and concordance of SCNAs among samples from the same individual.

#### Panel of normals creation

##### Small panel

We sequenced cell-free DNA (cfDNA) collected from blood plasma and genomic DNA (gDNA) collected from PBMCs from five healthy dogs. We generated a panel of normals (PoN) using both cfDNA and gDNA samples and found that the gDNA PoN showed less noise (lower MAD) than the cfDNA PoN. For this reason, we used the gDNA PoN for subsequent analysis with ichorCNA. We realigned these bam files to CanFam4, and this small panel was used as the panel of normals for files aligned to CanFam4. *<u>Large panel</u>*: We also created a panel of normals using previously-published normal whole-genome sequencing data from 92 dogs^54,91^, along with the five PBMC samples included in the small panel (Table SPON). This was used as the panel of normals for samples aligned to CanFam3.1.

#### Canine whole exome capture design

The exome capture panel was designed by the Vertebrate Genomics group at the Broad Institute, together with the Broad Genomics Platform, who provided input on the regions targeted in the Broad Human Somatic Exome Capture. The Bioinformatics team at Twist Bioscience then designed the baits to capture the targeted regions. This panel targets 20,257 genes derived from the Ensembl canine gene annotation Release 99, human cancer-specific targets (translated from hg19 to CanFam3.1 using liftOver^92^), and selected gene promoter sites and UTRs. The canine exome design targets approximately 36.6 Mb with 40.5 Mb of probe territory^55^ and is available as a product from Twist Bioscience.

#### Whole exome hybrid capture and sequencing

Pre-existing ULPWGS libraries were quantified and pooled prior to selection. The library pool along with the Canine Exome Capture v2 probe set were used as input into the xGEn Hyb and Wash kit (IDT). Post capture, libraries were sequenced on the Illumina HiSeq X platform using a 2x151 bp run, to a target depth of 150x coverage, achieving a mean target coverage of 131x.

#### Alignment and preprocessing of whole-exome sequencing data

##### Broad Genomics Platform and Blood biopsy group

ULPWGS sequencing data was preprocessed following the GATK best practices. Sequencing data was aligned to the CanFam3.1 reference genome^86^ using bwa^87^ version 0.7.15-r1140, and duplicate reads were flagged using Picard Tools MarkDuplicates. A VCF of known canid germline variation (34,191,821 SNPs and 11,943,064 indels from 674 dogs and other canids^91^) was used as the germline reference for BQSR. Preprocessing was performed using the “processing-for-variant-discovery” pipeline, with gatk version 4.1.8.1 and genomes-in-the-cloud version 2.3.1-1512499786.

### Simple somatic mutation calling

Simple somatic variant calling was performed on whole genome sequencing (WGS) data from 24 osteosarcoma and paired normal samples, as well as whole exome sequencing (WES) data from paired cfDNA samples from six of the same dogs. WGS data was subset to exome regions using the Samtools^93,94^ view command with four threads allocated and bam as the output file format. Data was preprocessed in the Terra Cloud Platform using the “Processing-for-variant-discovery” showcase workflow. To ensure the robustness of the somatic mutation calls, we used three different callers: Mutect2 v4.1.8.1, Strelka2 v2.9.10, and Octopus v0.7.4. Each tumor or cfDNA sample was run with its appropriate matched normal sample, and mutations were required to be called by two or more of the tools to be retained for downstream analysis. We used a VCF of germline variants in 674 dogs as a population variant resource. Mutect2 was run with the additional filter “minimum median read length” of 10 and the additional arguments: 20 downsample stride, 6 max reads per alignment start, and 6 max suspicious reads per alignment start. Tumor-cfDNA pair WLOS_0015 was excluded from subsequent concordance analysis because we detected only a single mutation in the tumor. This sample had a higher proportion of mutational calls flagged as potential cross-sample contamination by Mutect2 (70%; the other five tumor samples had a range of 26% - 40%), which may have contributed to the low number of variants passing filtration.

### Consensus of somatic mutations analysis

Raw mutation calls from each tool were filtered to include only “passing” mutations using Bcftools view function. Consensus mutation call sets from the three tools were created using Bcftools isec. SnpEff was used to annotate the calls and SnpSift was used to extract annotations into tab-delimited format. The intersection of the consensus mutation calls for each tumor - cfDNA pair was obtained using Bcftools isec.

## Data Availability

The raw sequence read data for all samples will be available in the NCBI Sequence Read Archive (SRA) and Integrated Canine Data Commons (ICDC) upon publication.

## Acknowledgements

Funding: This work was supported by the following grants from the National Institutes of Health: R01CA255319 (Karlsson, London), R37CA218570 (Karlsson, London), NIH U01CA224153 (London), NIH U01CA224182 (London). KM was supported by NCI fellowship F32CA247088. HLG is supported by K01OD028268-01A1. The content of this manuscript is solely the responsibility of the authors and does not necessarily represent the official views of the National Institutes of Health. Support was also provided by an American Cancer Society Research Scholar Grant (Gardner, Karlsson), an MIB Agents Outsmarting Osteosarcoma (Gardner, London), and the Dr. Eileen L. Berman and Stanley I. Berman Foundation. The authors also acknowledge the generous support of the Gerstner Family Foundation.

The authors would like to thank Paul Maza, DVM, PhD, Senior Lecturer in Anatomy at the Cornell University College of Veterinary Medicine, for guidance on the blood draw site diagrams and circulatory distance measurements. We would also like to thank Leslie Gaffney of the Broad Institute Communications Laboratory for her work designing the overview figure.

## Author contributions

**KM:** conceptualization, methodology, investigation, formal analysis, visualization, data curation, writing–original draft preparation, project administration, writing–review and editing, supervision

**CH:** methodology, formal analysis, data curation, writing–original draft preparation, writing–review and editing

**JR:** software, methodology, investigation, writing–original draft preparation, writing–review and editing

**MEW:** methodology, investigation, formal analysis, data curation,writing–review and editing

**DG:** methodology, investigation, writing–review and editing

**FLC:** methodology, data curation, writing–review and editing

**KX:** methodology, investigation, formal analysis, writing–original draft preparation, writing–review and editing

**EJK:** formal analysis, writing–review and editing

**RS:** methodology

**CP:** conceptualization, writing–review and editing

**VA:** conceptualization, methodology, resources, writing–review and editing, supervision

**CAL:** conceptualization, resources, investigation, data curation, project administration, writing–review and editing, supervision

**HLG:** conceptualization, methodology, resources, investigation, data curation, project administration, writing– review and editing, supervision

**EKK:** conceptualization, investigation, visualization, data curation, project administration, writing–review and editing, supervision

## Abbreviations

cfDNA: cell-free DNA
ng: nanograms
mL: milliliters
μL: microliters
bp: basepairs
Mb: megabases
PCR: polymerase chain reaction
ULPWGS: ultra low-pass whole-genome sequencing
WGS: whole-genome sequencing

## SUPPLEMENTARY FIGURE LEGENDS

**Figure SDOGS.** Overall the cohort of dogs enrolled with cancer was not significantly different from the healthy cohort for **(A)** sex, **(B)** spay/neuter status, **(C)** weight, and **(D)** breed ancestry. However, **(E)** dogs with cancer were significantly older than healthy dogs.

**Figure SMETRICS.** Comparison of liquid biopsy results between dogs and humans by cancer type. **(A)** ANOVA results showing that both species and cancer type significantly affected cfDNA concentration and fragment size ratio, while only cancer type had an effect on tumor fraction. The species difference in fragment size ratio persisted even when limiting dog samples to EDTA only. Boxplots display distribution by species and cancer type of **(B)** tumor fraction, **(C)** fragment size ratio, and **(D)** cfDNA concentration. **(E)** While all dog cancers and human cancers had significantly different mean tumor fractions, removing the canine lymphomas nearly eliminates the effect.

**Figure SREPLICATES.** Plots of technical replicate values from two DNA extractions of the same blood draw. **(A)** cfDNA concentration, **(B)** Tumor fraction, **(C)** fragment size ratio. All are highly concordant, with tumor fraction showing the least variability (R=0.99) and fragment size ratio showing the most variability (R=0.76).

**Figure SINPUT.** Plots of replicate libraries constructed using different input amounts of the same DNA samples. **(A)** Tumor fraction. **(B)** Fragment size ratio. Again, fragment size was more variable than tumor fraction.

**Figure SCOMPARE.** Correlation of the three cfDNA metrics between healthy dogs, dogs with lymphoma, and dogs with other cancers, visualized as **(A)** forest plots, and scatter plots**. (B)** cfDNA concentration was strongly positively correlated with tumor fraction in lymphomas only. **(C)** cfDNA concentration was significantly negatively correlated with fragment size ratio in both lymphomas and healthy dogs, but not other cancers. **(D)** Tumor fraction was positively correlated with fragment size ratio in other cancer types, but negatively correlated in lymphomas.

**Figure SSPEAR.** Spearman correlations showing the concordance between log2 read depth ratios of tumor and cfDNA in 24 dogs with paired osteosarcoma tumor and liquid biopsy samples.

**Figure SCONCORD.** Concordance of simple somatic mutation calls between tumor and cfDNA sequencing in five dogs with osteosarcoma. **(A)** Overall, we detected significantly more mutations in the cfDNA than in the tumor. Consistent with this, **(B)** somatic mutations identified in the cfDNA were less likely to be identified in the tumor, **(C)** a higher proportion of the mutations identified in an individual dog were identified in the cfDNA, **(D)** median allele fractions were lower in the cfDNA than in the tumor, and **(E)** heterogeneity (MATH) scores were higher in the cfDNA than in the tumor.

**Figure_SSWAP. Pairwise** percent concordance of genotypes over 9,137 common SNP sites between all tumor and cfDNA samples in the simple somatic mutation concordance analysis. No evidence of sample swap is identified.

**Figure SBREEDMETRICS.** Overview of breed statistics in the study cohort. **(A)** Thirty-eight breeds were represented by at least one dog. The most commonly represented breeds were labrador retrievers (n=22) and golden retrievers (n=17). **(B)** cfDNA concentration was significantly different between labrador retrievers and other dogs. **(C)** Removing the lymphomas from the analysis eliminates this difference. There are no significant differences between labrador retrievers, golden retrievers, and other dogs in **(D)** tumor fraction or **(E)** tumor fraction excluding lymphomas, or in **(F)** fragment size ratios with or **(G)** without lymphomas.

**Figure STUBES.** Blood collection tube type had an effect on cfDNA metrics. **(A)** cfDNA concentration was significantly higher in samples collected in EDTA vs Streck tubes from dogs with non-lymphoma cancers, suggesting that more white blood cell lysis was occurring in the EDTA samples. There were no significant differences in **(B)** tumor fraction or **(C)** fragment size ratio between samples collected in EDTA and Streck tubes, including in five dogs where a sample was collected in both tube types. Samples from dogs with lymphoma collected in Streck tubes were significantly higher in cfDNA concentration and tumor fraction and lower in fragment size ratio than samples from dogs with other cancer types collected in Streck tubes.

**Figure SDRAW. (A)** Tumor fraction was significantly associated with circulation distance between the tumor and blood draw site. **(B)** Fragment size ratio was significantly associated with circulation distance between the tumor and blood draw site. **(C)** cfDNA concentration was not significantly associated with circulation distance between the tumor and blood draw site.

**Figure STIME.** In dogs where longitudinal liquid biopsies were taken 2-3.5 hours apart, time of day had no significant effect on **(A)** cfDNA concentration, **(B)** tumor fraction or **(C)** fragment size ratio. There was also no significant effect of morning blood draw time vs afternoon on the three metrics **(D-F)**.

**Figure SSHORT.** cfDNA metrics from the acute response monitoring study of four dogs with lymphoma pre- and post-treatment with L-asparaginase. Pink bar denotes treatment time point. **(A)** Lymph node sizes for the three dogs with DLBCL. **(B)** cfDNA concentration, **(C)** tumor fraction or **(D)** fragment size ratio. **(E)** Total lymph node sizes by cfDNA concentration, tumor fraction, or fragment size ratio for the three dogs with DLBCL.

**Figure SLONBOX.** Multiple samples from dogs with lymphoma or osteosarcoma over the course of treatment. Plots compare initial pre-treatment of pre-amputation time points (“disease present”), remission or post- amputation time points (“no disease”), and “progressive disease” time points. **(A)** cfDNA concentration was associated with disease status in dogs with lymphoma. **(B)** Tumor fraction was significantly associated with disease status in lymphomas, and between “disease present” and “no disease” in osteosarcomas. **(C)** Fragment size ratio was significantly associated with disease status in osteosarcomas but not lymphomas.

**Figure SLONGDOG_LSA_CONC.** Longitudinal cfDNA concentrations over the course of treatment for 34 dogs with DLBCL. Pink bar denotes progressive disease time point.

**Figure SLONGDOG_LSA_TF.** Longitudinal tumor fractions over the course of treatment for 34 dogs with DLBCL. Pink bar denotes progressive disease time point.

**Figure SLONGDOG_LSA_FRAG.** Longitudinal fragment size ratios over the course of treatment for 34 dogs with DLBCL. Pink bar denotes progressive disease time point.

**Figure SLONGDOG_OS_TF.** Longitudinal tumor fractions over the course of treatment for 21 dogs with OS. Pink bar denotes progressive disease time point.

**Figure SLONGDOG_OS_FRAG.** Longitudinal fragment size ratios over the course of treatment for 21 dogs with OS. Pink bar denotes progressive disease time point.

**Figure SFOREST.** Forest plot of Cox Proportional Hazards model of (A) cfDNA concentration, tumor fraction, and fragment size ratio at day 0 in dogs with lymphoma with longitudinal data. cfDNA concentration was significantly associated with PFS. **(B)** cfDNA concentration, tumor fraction, fragment size ratio, and disease status at day 7 in dogs with lymphoma with longitudinal data. Tumor fraction was significantly associated with PFS.

## References

1. Chu, D. & Park, B. H. Liquid biopsy: unlocking the potentials of cell-free DNA. Virchows Arch. 471, 147– 154 (2017).

2. FDA Approves Blood Tests That Can Help Guide Cancer Treatment. National Cancer Institute https://www.cancer.gov/news-events/cancer-currents-blog/2020/fda-guardant-360-foundation-one-cancer-liquid-biopsy (2020).

3. Kwapisz, D. The first liquid biopsy test approved. Is it a new era of mutation testing for non-small cell lung cancer? Ann Transl Med 5, 46 (2017).

4. Woodhouse, R. et al. Clinical and analytical validation of FoundationOne Liquid CDx, a novel 324-Gene cfDNA-based comprehensive genomic profiling assay for cancers of solid tumor origin. PLoS One 15, e0237802 (2020).

5. Bauml, J. M. et al. Clinical validation of Guardant360 CDx as a blood-based companion diagnostic for sotorasib. Lung Cancer 166, 270–278 (2022).

6. Christou, N. et al. Circulating Tumour Cells, Circulating Tumour DNA and Circulating Tumour miRNA in Blood Assays in the Different Steps of Colorectal Cancer Management, a Review of the Evidence in 2019. BioMed Research International vol. 2019 1–11 Preprint at 10.1155/2019/5953036 (2019).

7. Manier, S. et al. Whole-exome sequencing of cell-free DNA and circulating tumor cells in multiple myeloma. Nat. Commun. 9, 1691 (2018).

8. Parikh, A. R. et al. Liquid versus tissue biopsy for detecting acquired resistance and tumor heterogeneity in gastrointestinal cancers. Nat. Med. 25, 1415–1421 (2019).

9. Stover, D. G. et al. Association of Cell-Free DNA Tumor Fraction and Somatic Copy Number Alterations With Survival in Metastatic Triple-Negative Breast Cancer. J. Clin. Oncol. 36, 543–553 (2018).

10. Adalsteinsson, V. A. et al. Scalable whole-exome sequencing of cell-free DNA reveals high concordance with metastatic tumors. Nat. Commun. 8, 1–13 (2017).

11. Thompson, J. C. et al. Detection of Therapeutically Targetable Driver and Resistance Mutations in Lung Cancer Patients by Next-Generation Sequencing of Cell-Free Circulating Tumor DNA. Clin. Cancer Res. 22, 5772–5782 (2016).

12. Lockwood, C. M. et al. Recommendations for Cell-Free DNA Assay Validations: A Joint Consensus Recommendation of the Association for Molecular Pathology and College of American Pathologists. J. Mol. Diagn. 25, 876–897 (2023).

13. Heitzer, E., Haque, I. S., Roberts, C. E. S. & Speicher, M. R. Current and future perspectives of liquid biopsies in genomics-driven oncology. Nat. Rev. Genet. 20, 71–88 (2019).

14. Kang, Q. et al. Comparative analysis of circulating tumor DNA stability In K3EDTA, Streck, and CellSave blood collection tubes. Clin. Biochem. 49, 1354–1360 (2016).

15. Bettegowda, C. et al. Detection of circulating tumor DNA in early- and late-stage human malignancies. Sci. Transl. Med. 6, 224ra24 (2014).

16. Tóth, K. et al. Circadian Rhythm of Methylated Septin 9, Cell-Free DNA Amount and Tumor Markers in Colorectal Cancer Patients. Pathol. Oncol. Res. 23, 699–706 (2017).

17. Madsen, A. T., Hojbjerg, J. A., Sorensen, B. S. & Winther-Larsen, A. Day-to-day and within-day biological variation of cell-free DNA. EBioMedicine 49, 284–290 (2019).

18. Atamaniuk, J. et al. Increased concentrations of cell-free plasma DNA after exhaustive exercise. Clin. Chem. 50, 1668–1670 (2004).

19. Mavropalias, G. et al. Changes in plasma hydroxyproline and plasma cell-free DNA concentrations after higher- versus lower-intensity eccentric cycling. Eur. J. Appl. Physiol. 121, 1087–1097 (2021).

20. Meddeb, R., Pisareva, E. & Thierry, A. R. Guidelines for the Preanalytical Conditions for Analyzing Circulating Cell-Free DNA. Clin. Chem. 65, 623–633 (2019).

21. Ignatiadis, M., Sledge, G. W. & Jeffrey, S. S. Liquid biopsy enters the clinic - implementation issues and future challenges. Nat. Rev. Clin. Oncol. 18, 297–312 (2021).

22. Cisneros-Villanueva, M. et al. Cell-free DNA analysis in current cancer clinical trials: a review. Br. J. Cancer 126, 391–400 (2022).

23. Ungerer, V., Bronkhorst, A. J. & Holdenrieder, S. Preanalytical variables that affect the outcome of cell-free DNA measurements. Crit. Rev. Clin. Lab. Sci. 57, 484–507 (2020).

24. WHO Guidelines on Drawing Blood: Best Practices in Phlebotomy. (World Health Organization, Geneva, 2010).

25. Jiang, C. Y. et al. It’s not ‘just a tube of blood’: principles of protocol development, sample collection, staffing and budget considerations for blood-based biomarkers in immunotherapy studies. J Immunother Cancer 9, (2021).

26. Cheon, D.-J. & Orsulic, S. Mouse models of cancer. Annu. Rev. Pathol. 6, 95–119 (2011).

27. Parasuraman, S., Raveendran, R. & Kesavan, R. Blood sample collection in small laboratory animals. J. Pharmacol. Pharmacother. 1, 87–93 (2010).

28. Guidelines for Survival Blood Collection in Mice and Rats. https://oacu.oir.nih.gov/system/files/media/file/2022-12/b2-Survival_Blood_Collection_Mice_Rats.pdf (2023).

29. Gardner, H. L., Fenger, J. M. & London, C. A. Dogs as a Model for Cancer. Annu Rev Anim Biosci 4, 199– 222 (2016).

30. Nationwide releases findings on cancer in dogs. American Veterinary Medical Association https://www.avma.org/news/nationwide-releases-findings-cancer-dogs.

31. Veterinary Research & Analytics Information. Nationwide Pet Insurance https://www.petinsurance.com/veterinarians/research/ (2019).

32. LeBlanc, A. K. et al. Perspectives from man’s best friend: National Academy of Medicine’s Workshop on Comparative Oncology. Sci. Transl. Med. 8, 324ps5 (2016).

33. Paoloni, M. & Khanna, C. Translation of new cancer treatments from pet dogs to humans. Nat. Rev. Cancer 8, 147–156 (2008).

34. Schaefer, D. M. W. et al. Quantification of plasma DNA as a prognostic indicator in canine lymphoid neoplasia. Vet. Comp. Oncol. 5, 145–155 (2007).

35. Gelaleti, G. B. et al. Short interspersed CAN SINE elements as prognostic markers in canine mammary neoplasia. Oncol. Rep. 31, 435–441 (2014).

36. Beck, J. et al. Genome aberrations in canine mammary carcinomas and their detection in cell-free plasma DNA. PLoS One 8, e75485 (2013).

37. Beffagna, G. et al. Circulating Cell-Free DNA in Dogs with Mammary Tumors: Short and Long Fragments and Integrity Index. PLoS One 12, e0169454 (2017).

38. Lee, K.-H., Shin, T.-J., Kim, W.-H. & Cho, J.-Y. Author Correction: Methylation of LINE-1 in cell-free DNA serves as a liquid biopsy biomarker for human breast cancers and dog mammary tumors. Sci. Rep. 9, 17459 (2019).

39. Tagawa, M., Shimbo, G., Inokuma, H. & Miyahara, K. Quantification of plasma cell-free DNA levels in dogs with various tumors. Journal of Veterinary Diagnostic Investigation vol. 31 836–843 Preprint at 10.1177/1040638719880245 (2019).

40. Lorch, G. et al. Identification of Recurrent Activating HER2 Mutations in Primary Canine Pulmonary Adenocarcinoma. Clin. Cancer Res. 25, 5866–5877 (2019).

41. Goggs, R. Effect of sample type on plasma concentrations of cell-free DNA and nucleosomes in dogs. Vet. Rec. Open 6, e000357 (2019).

42. Schabort, J. J. et al. ANK2 Hypermethylation in Canine Mammary Tumors and Human Breast Cancer. Int. J. Mol. Sci. 21, (2020).

43. Favaro, P. F. et al. Feasibility and promise of circulating tumor DNA analysis in dogs with naturally-occurring sarcoma. bioRxiv 2020.08.20.260349 (2020) doi:10.1101/2020.08.20.260349.

44. Tagawa, M. et al. Quantitative analysis of the BRAF V595E mutation in plasma cell-free DNA from dogs with urothelial carcinoma. PLoS One 15, e0232365 (2020).

45. Prouteau, A. et al. Circulating tumor DNA is detectable in canine histiocytic sarcoma, oral malignant melanoma, and multicentric lymphoma. Sci. Rep. 11, 877 (2021).

46. Kruglyak, K. M. et al. Blood-based liquid biopsy for comprehensive cancer genomic profiling using next-generation sequencing: An emerging paradigm for non-invasive cancer detection and management in dogs. Front. Vet. Sci. 8, 704835 (2021).

47. Flory, A. et al. Clinical validation of a next-generation sequencing-based multi-cancer early detection ‘liquid biopsy’ blood test in over 1,000 dogs using an independent testing set: The CANcer Detection in Dogs (CANDiD) study. PLoS One 17, e0266623 (2022).

48. Kim, J. et al. Cell-Free DNA as a Diagnostic and Prognostic Biomarker in Dogs With Tumors. Front Vet Sci 8, 735682 (2021).

49. Guil-Luna, S. et al. Analysis of cell-free DNA concentration, fragmentation patterns and TP53 gene expression in mammary tumor-bearing dogs: A pilot study. Front Vet Sci 10, 1157878 (2023).

50. Cristiano, S. et al. Genome-wide cell-free DNA fragmentation in patients with cancer. Nature 570, 385– 389 (2019).

51. Peneder, P. et al. Multimodal analysis of cell-free DNA whole-genome sequencing for pediatric cancers with low mutational burden. Nat. Commun. 12, 3230 (2021).

52. Rickles-Young, M. et al. Assay Validation of Cell-Free DNA Shallow Whole-Genome Sequencing to Determine Tumor Fraction in Advanced Cancers. J. Mol. Diagn. 26, 413–422 (2024).

53. Cescon, D. W., Bratman, S. V., Chan, S. M. & Siu, L. L. Circulating tumor DNA and liquid biopsy in oncology. Nat Cancer 1, 276–290 (2020).

54. Gardner, H. L. et al. Canine osteosarcoma genome sequencing identifies recurrent mutations in DMD and the histone methyltransferase gene SETD2. Commun Biol 2, 266 (2019).

55. Kate Megquier Ross Swofford Holly Corbitt Tera Bowers Justin Abreu Sheli McDonough Micah Rickles- Young Anna Koutoulas Junko Tsuji Edyta Malolepsza Carrie Cibulskis Brendan Blumenstiel Keith McKenna Scott McCuine Paul Frere Patrick Boyle Frances L. Chen Heather L. Gardner Cheryl A. London Elinor K. Karlsson. Broad-UMass-Chan-Canine-Exome-Whitepaper-2023. https://karlssonlab.org/2024/03/27/exome/broad-umass-chan-canine-exome-whitepaper-2023/.

56. Benjamin, D. et al. Calling Somatic SNVs and Indels with Mutect2. bioRxiv 861054 (2019) doi:10.1101/861054.

57. Kim, S. et al. Strelka2: fast and accurate calling of germline and somatic variants. Nat. Methods 15, 591– 594 (2018).

58. Cooke, D. P., Wedge, D. C. & Lunter, G. A unified haplotype-based method for accurate and comprehensive variant calling. Nat. Biotechnol. 39, 885–892 (2021).

59. Wang, P. P. S., Parker, W. T., Branford, S. & Schreiber, A. W. BAM-matcher: a tool for rapid NGS sample matching. Bioinformatics 32, 2699–2701 (2016).

60. Pereira, B. et al. Cell-free DNA captures tumor heterogeneity and driver alterations in rapid autopsies with pre-treated metastatic cancer. Nat. Commun. 12, 3199 (2021).

61. Wu, X., Song, P., Guo, L., Ying, J. & Li, W. Mutant-Allele Tumor Heterogeneity, a Favorable Biomarker to Assess Intra-Tumor Heterogeneity, in Advanced Lung Adenocarcinoma. Front. Oncol. 12, 888951 (2022).

62. Hsiehchen, D., Espinoza, M., Gerber, D. E. & Beg, M. S. Clinical and biological determinants of circulating tumor DNA detection and prognostication using a next-generation sequencing panel assay. Cancer Biol. Ther. 22, 455–464 (2021).

63. Lui, Y. Y. N. et al. Predominant hematopoietic origin of cell-free DNA in plasma and serum after sex-mismatched bone marrow transplantation. Clin. Chem. 48, 421–427 (2002).

64. Razavi, P. et al. High-intensity sequencing reveals the sources of plasma circulating cell-free DNA variants. Nat. Med. 25, 1928–1937 (2019).

65. Damascelli, B., Tichà, V., Repetti, E. & Dorji, T. Beyond Standard Practice in Liquid Biopsy: Selective Venous Sampling. J. Vasc. Interv. Radiol. 32, 668–671 (2021).

66. Wagner, J. T. et al. Diurnal stability of cell-free DNA and cell-free RNA in human plasma samples. Sci. Rep. 10, 16456 (2020).

67. Korabecna, M., Horinek, A., Bila, N. & Opatrna, S. Circadian Rhythmicity and Clearance of Cell-Free DNA in Human Plasma. in Circulating Nucleic Acids in Plasma and Serum 195–198 (Springer Netherlands, 2011).

68. Feriel, J., Tchipeva, D. & Depasse, F. Effects of circadian variation, lifestyle and environment on hematological parameters: A narrative review. Int. J. Lab. Hematol. 43, 917–926 (2021).

69. Busza, A. et al. 0028 Systematic Review: Time of day differences in complete blood count values. Sleep 46, A12–A12 (2023).

70. Zhu, X. et al. In vivo flow cytometry reveals a circadian rhythm of circulating tumor cells. Light Sci Appl 10, 110 (2021).

71. Spindler, K.-L. G. & Jakobsen, A. Circulating tumor DNA: Response Evaluation Criteria in Solid Tumors - can we RECIST? Focus on colorectal cancer. Ther. Adv. Med. Oncol. 15, 17588359231171580 (2023).

72. Watanabe, K., Nakamura, Y. & Low, S.-K. Clinical implementation and current advancement of blood liquid biopsy in cancer. J. Hum. Genet. 66, 909–926 (2021).

73. Erdi, Y. E. Limits of Tumor Detectability in Nuclear Medicine and PET. Mol. Imaging Radionucl. Ther. 21, 23–28 (2012).

74. Eisenhauer, E. A. et al. New response evaluation criteria in solid tumours: revised RECIST guideline (version 1.1). Eur. J. Cancer 45, 228–247 (2009).

75. Terasawa, H. et al. Circulating tumor DNA dynamics analysis in a xenograft mouse model with esophageal squamous cell carcinoma. World J. Gastroenterol. 27, 7134–7143 (2021).

76. Walls, G. M. et al. Early circulating tumour DNA kinetics measured by ultra-deep next-generation sequencing during radical radiotherapy for non-small cell lung cancer: a feasibility study. Radiat. Oncol. 15, 132 (2020).

77. Diehl, F. et al. Circulating mutant DNA to assess tumor dynamics. Nat. Med. 14, 985–990 (2008).

78. Stadler, J.-C. et al. Current and Future Clinical Applications of ctDNA in Immuno-Oncology. Cancer Res. 82, 349–358 (2022).

79. Thompson, J. C., Scholes, D. G., Carpenter, E. L. & Aggarwal, C. Molecular response assessment using circulating tumor DNA (ctDNA) in advanced solid tumors. Br. J. Cancer 129, 1893–1902 (2023).

80. McCleary-Wheeler, A. L. et al. Next-generation sequencing-based liquid biopsy may be used for detection of residual disease and cancer recurrence monitoring in dogs. Am. J. Vet. Res. 1–8 (2023).

81. Mouliere, F. et al. Enhanced detection of circulating tumor DNA by fragment size analysis. Sci. Transl. Med. 10, (2018).

82. Lau, J. H. G., Drake, W. & Matson, M. The current role of venous sampling in the localization of endocrine disease. Cardiovasc. Intervent. Radiol. 30, 555–570 (2007).

83. Farrelly, C. et al. Correlation of Peripheral Vein Tumour Marker Levels, Internal Iliac Vein Tumour Marker Levels and Radical Prostatectomy Specimens in Patients with Prostate Cancer and Borderline High Prostate-Specific Antigen: A Pilot Study. Cardiovasc. Intervent. Radiol. 39, 724–731 (2016).

84. Denève, E. et al. Capture of viable circulating tumor cells in the liver of colorectal cancer patients. Clin. Chem. 59, 1384–1392 (2013).

85. Buscail, E. et al. Tumor-proximal liquid biopsy to improve diagnostic and prognostic performances of circulating tumor cells. Mol. Oncol. 13, 1811–1826 (2019).

86. Hoeppner, M. P. et al. An improved canine genome and a comprehensive catalogue of coding genes and non-coding transcripts. PLoS One 9, e91172 (2014).

87. Li, H. & Durbin, R. Fast and accurate short read alignment with Burrows-Wheeler transform. Bioinformatics 25, 1754–1760 (2009).

88. Wang, C. et al. A novel canine reference genome resolves genomic architecture and uncovers transcript complexity. Commun Biol 4, 185 (2021).

89. Ramírez, F. et al. deepTools2: a next generation web server for deep-sequencing data analysis. Nucleic Acids Res. 44, W160–5 (2016).

90. Lai, D. & Shah, S. HMMcopy: Copy number prediction with correction for GC and mappability bias for HTS data. R package version 1, (2012).

91. Morrill, K., et al. Ancestry-inclusive dog genomics challenges popular breed stereotypes. Science in press, (2022).

92. Hinrichs, A. S. et al. The UCSC Genome Browser Database: update 2006. Nucleic Acids Res. 34, D590–8 (2006).

93. Li, H. et al. The Sequence Alignment/Map format and SAMtools. Bioinformatics 25, 2078–2079 (2009).

94. Danecek, P. et al. Twelve years of SAMtools and BCFtools. Gigascience 10, (2021).

95. RDevelopment CORE TEAM, R. & Others. R: A language and environment for statistical computing. Preprint at (2023).

