## Supplementary Figures for "Impact of preanalytical factors on liquid biopsy in the canine cancer model"

### Figure SDOGS

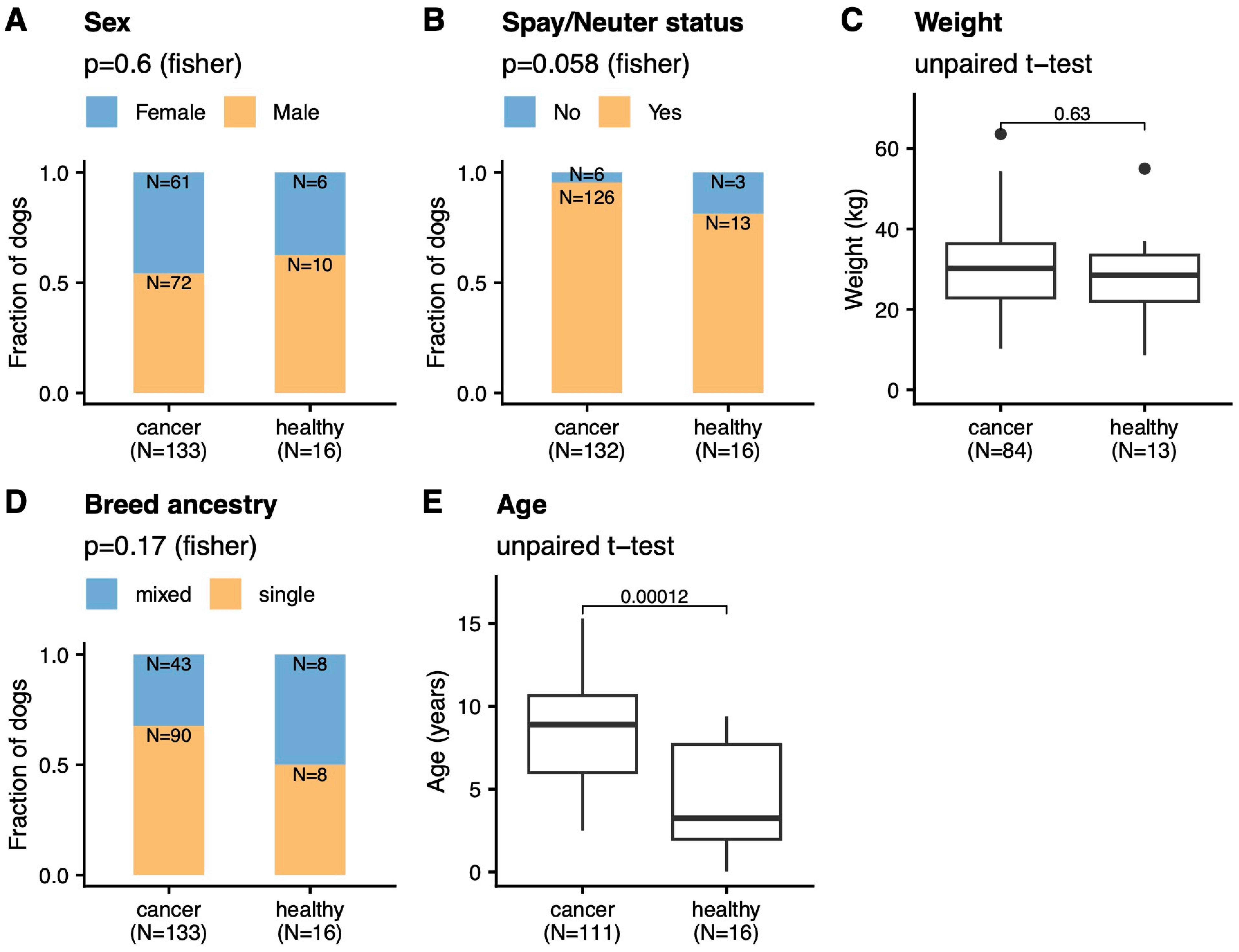

### Figure SMETRICS

#### A. Anova analysis on liquid biopsy metrics by species & cancer type (all dogs)

| statName | Effect | DFn | DFd | F | p | p<.05 | ges |
|---|---|---|---|---|---|---|---|
| Fragment size ratio | Cancer_type | 21 | 260 | 3.524 | 8.99e−07 | * | 0.222 |
| Fragment size ratio | species | 1 | 260 | 30.036 | 1e−07 | * | 0.104 |
| Tumor fraction | Cancer_type | 18 | 614 | 8.903 | 9.88e−22 | * | 0.207 |
| Tumor fraction | species | 1 | 614 | 0.121 | 0.728 | | 0.000 |
| cfDNA concentration | Cancer_type | 23 | 758 | 10.360 | 3.91e−32 | * | 0.239 |
| cfDNA concentration | species | 1 | 758 | 6.003 | 0.015 | * | 0.008 |

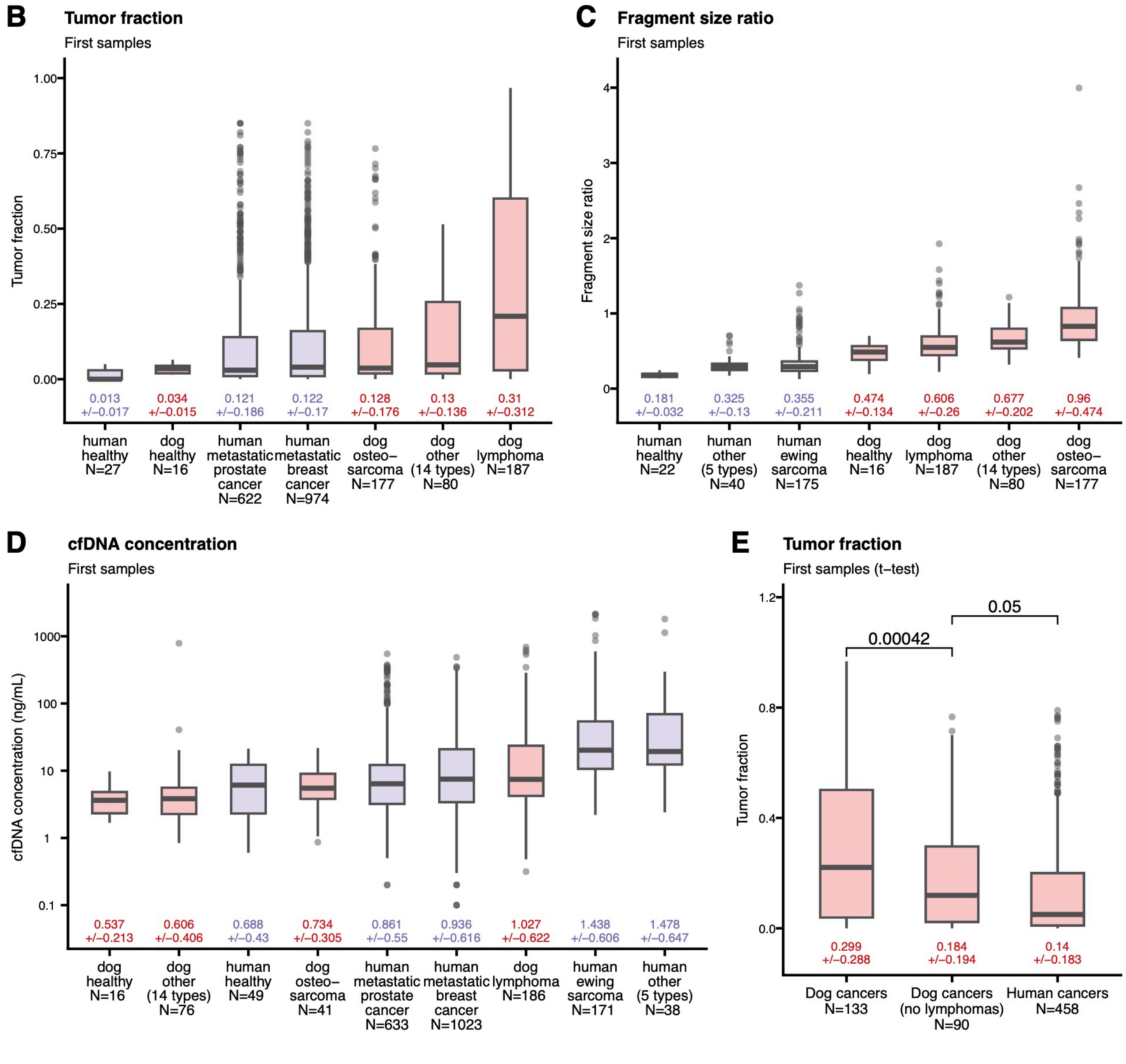

### Figure SREPLICATES

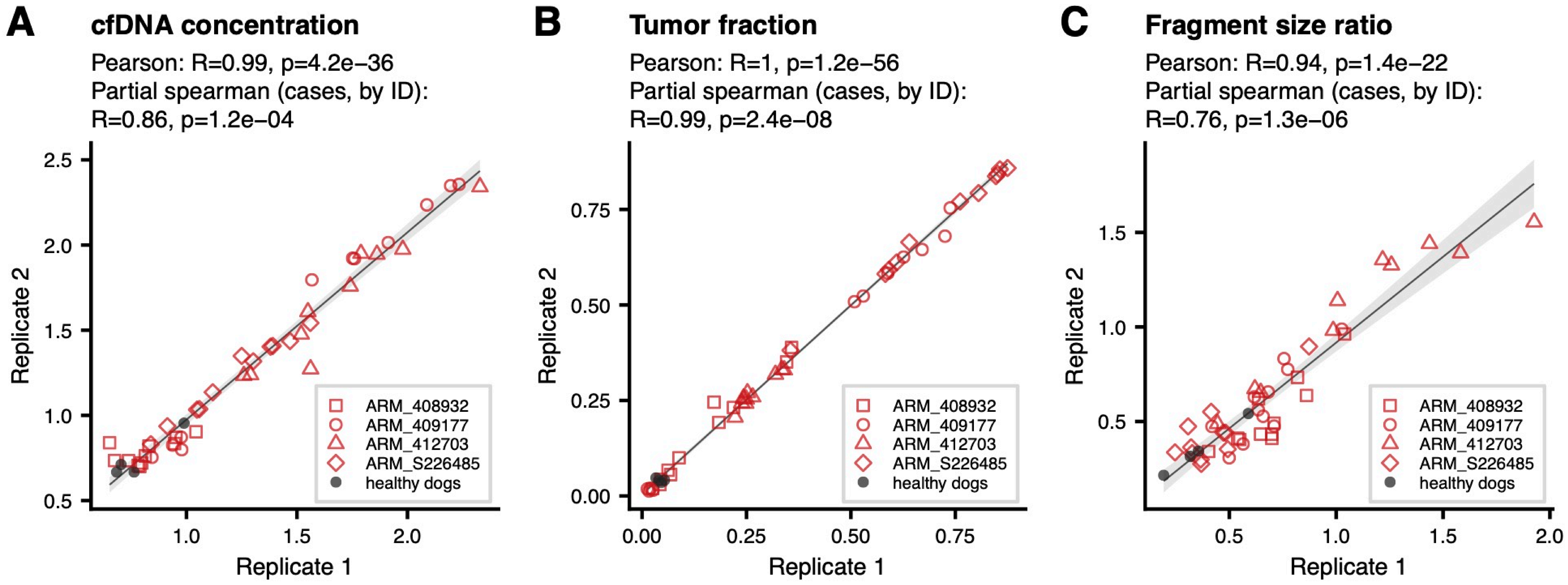

### Figure SINPUT

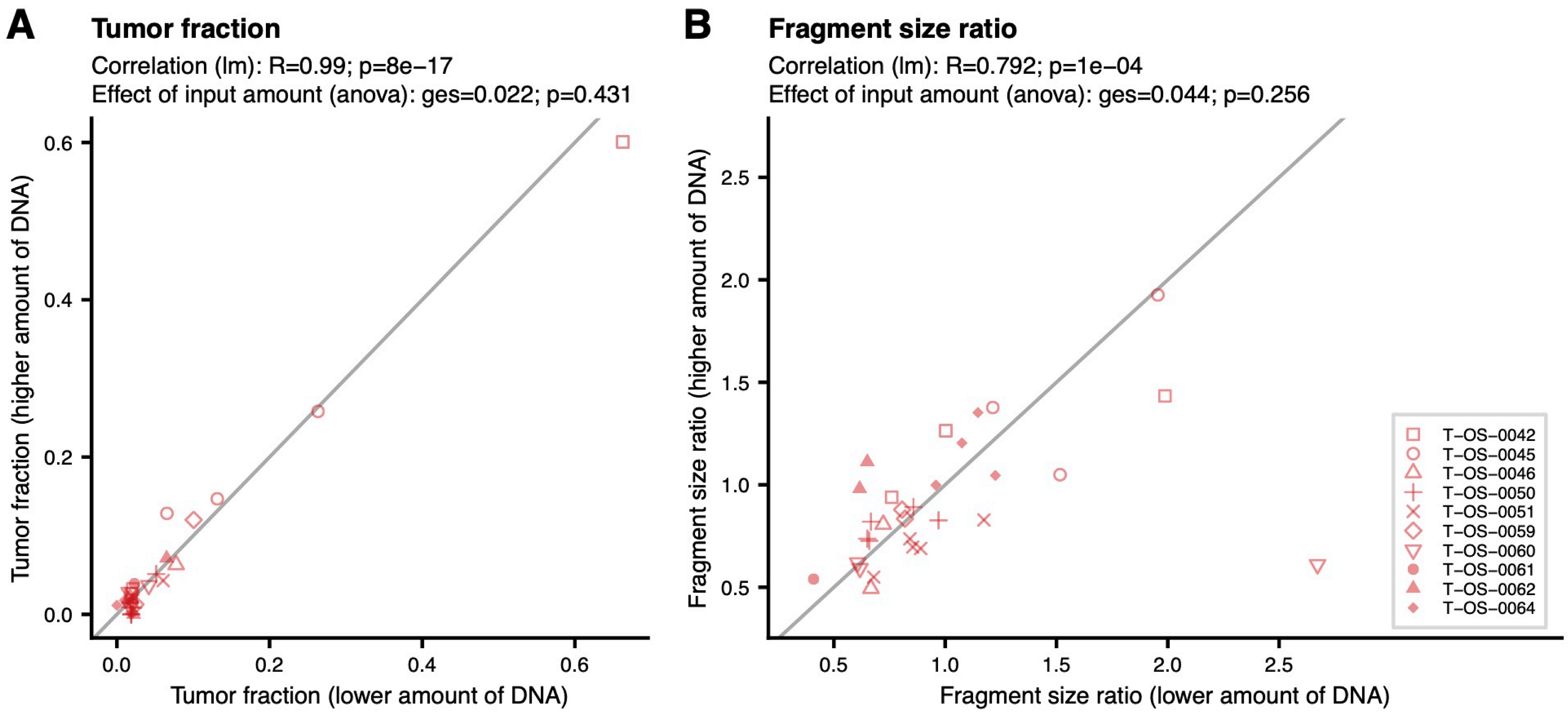

### Figure SCOMPARE

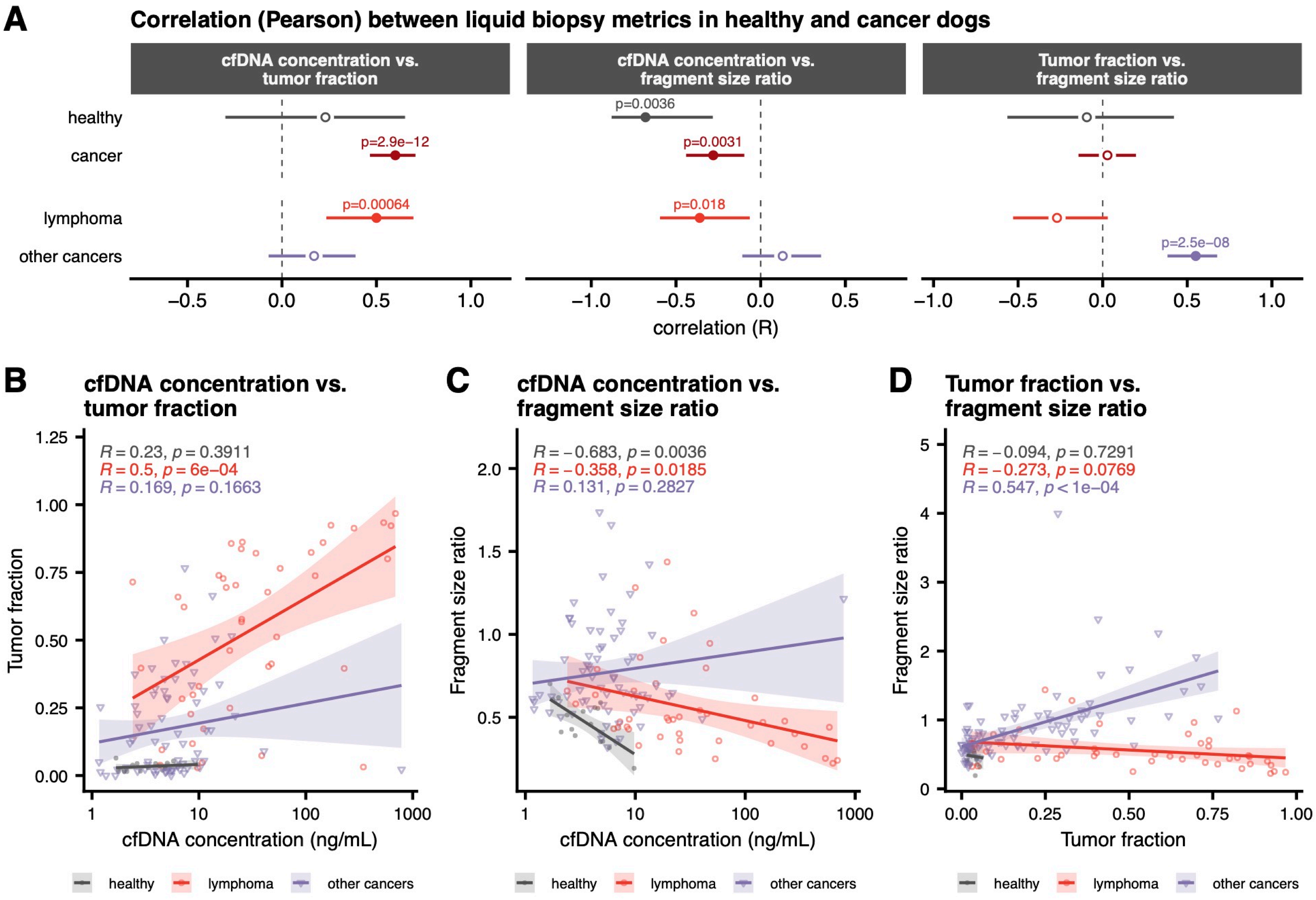

### Figure SSPEAR

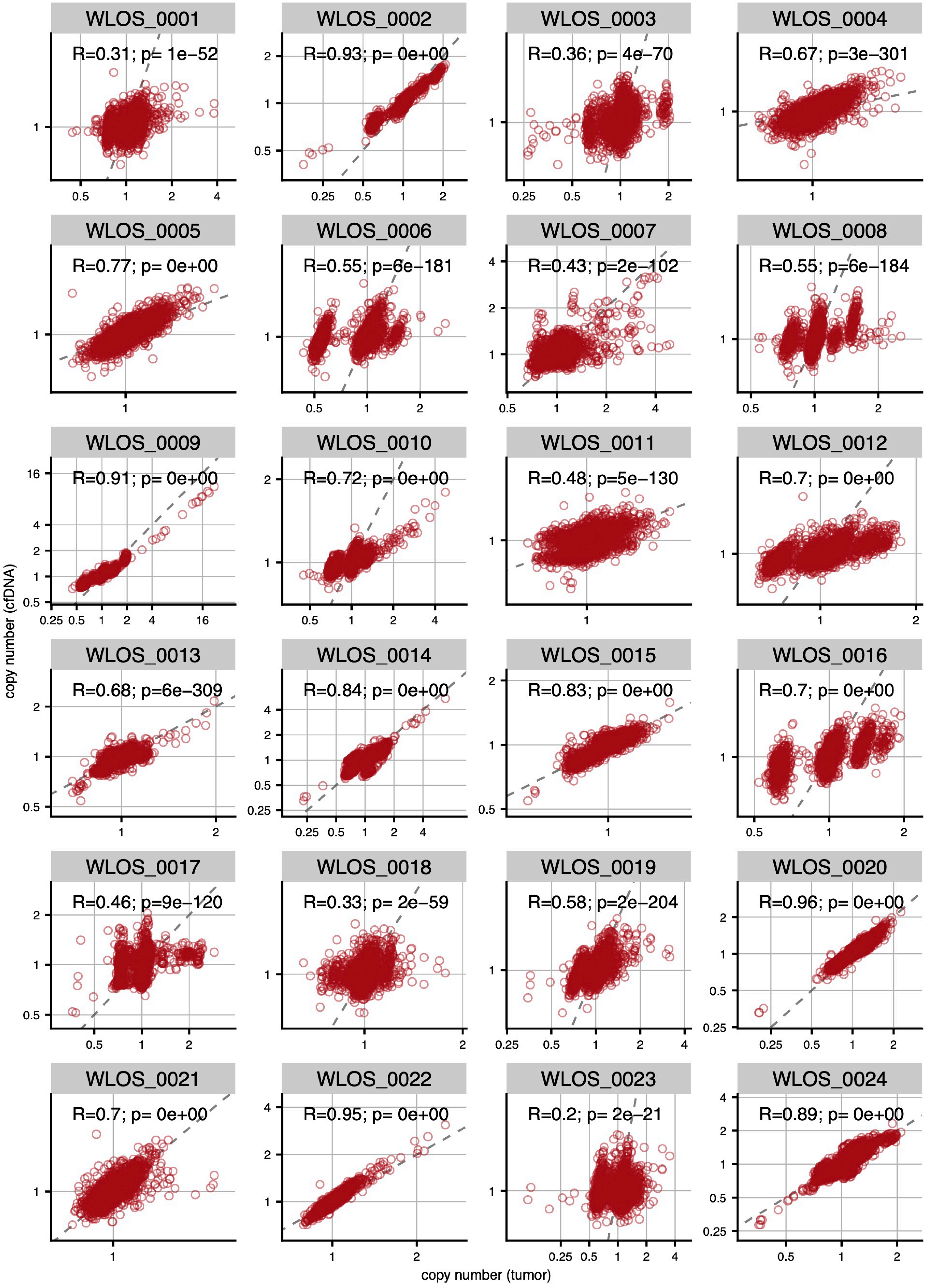

### Figure SCONCORD

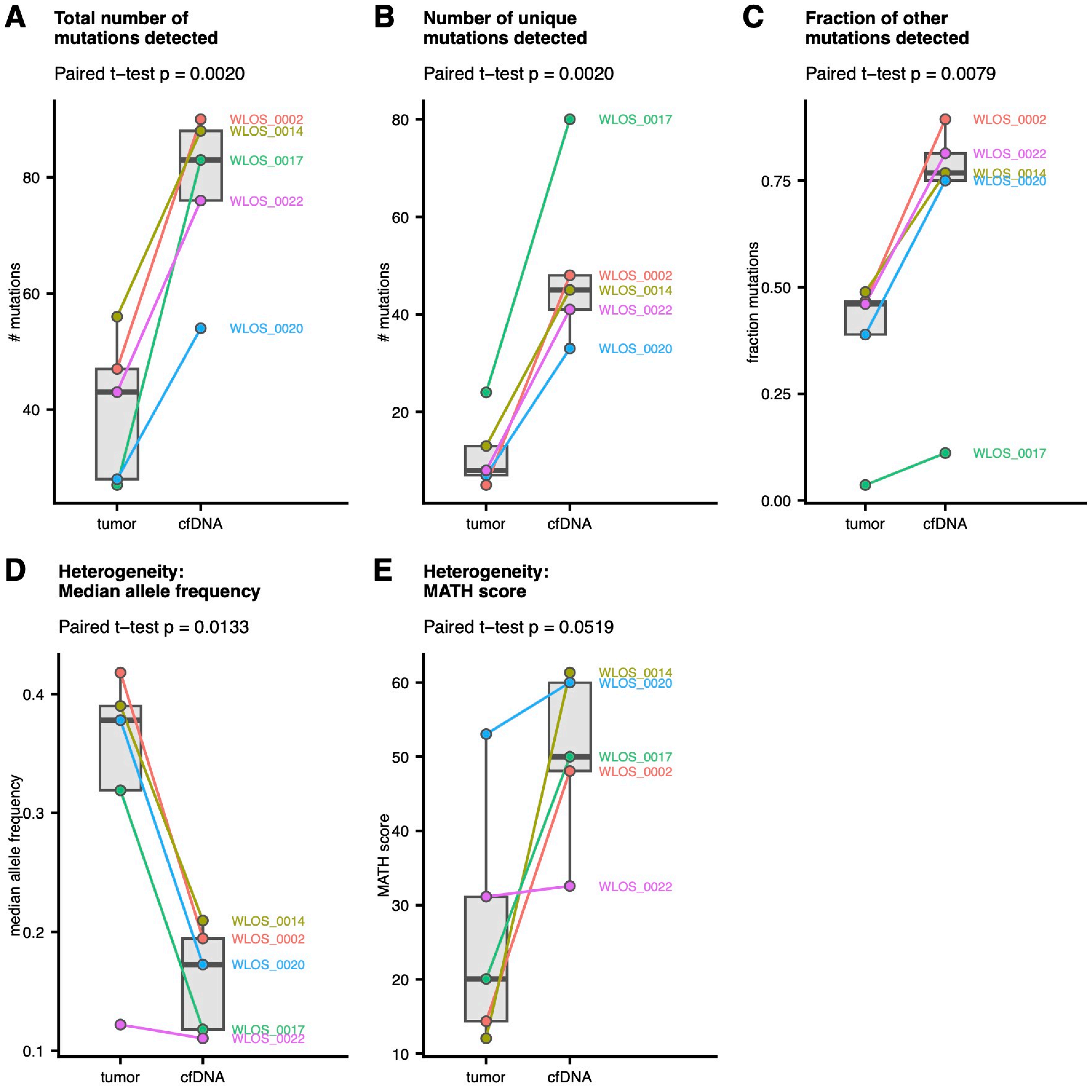

**Figure SSWAP**

| id | WLOS_0002_tumor | WLOS_0002_Plasma_1 | WLOS_0014_tumor | WLOS_0014_Plasma_1 | WLOS_0015_tumor | WLOS_0015_Plasma_1 | WLOS_0017_tumor | WLOS_0017_Plasma_1 | WLOS_0020_tumor | WLOS_0020_Plasma_1 | WLOS_0022_tumor | WLOS_0022_Plasma_1 | id |
|---|---|---|---|---|---|---|---|---|---|---|---|---|---|
| | 1.00 | 0.94 | 0.56 | 0.52 | 0.56 | 0.52 | 0.57 | 0.54 | 0.58 | 0.52 | 0.58 | 0.54 | WLOS_0002_tumor |
| | 0.94 | 1.00 | 0.52 | 0.55 | 0.52 | 0.56 | 0.53 | 0.57 | 0.54 | 0.57 | 0.54 | 0.58 | WLOS_0002_Plasma_1 |
| | 0.56 | 0.52 | 1.00 | 0.95 | 0.57 | 0.53 | 0.57 | 0.53 | 0.57 | 0.51 | 0.58 | 0.54 | WLOS_0014_tumor |
| | 0.52 | 0.55 | 0.95 | 1.00 | 0.53 | 0.56 | 0.53 | 0.57 | 0.53 | 0.56 | 0.54 | 0.58 | WLOS_0014_Plasma_1 |
| | 0.56 | 0.52 | 0.57 | 0.53 | 1.00 | 0.95 | 0.59 | 0.55 | 0.58 | 0.53 | 0.59 | 0.54 | WLOS_0015_tumor |
| | 0.52 | 0.56 | 0.53 | 0.56 | 0.95 | 1.00 | 0.54 | 0.58 | 0.54 | 0.57 | 0.54 | 0.58 | WLOS_0015_Plasma_1 |
| | 0.57 | 0.53 | 0.57 | 0.53 | 0.59 | 0.54 | 1.00 | 0.95 | 0.60 | 0.55 | 0.59 | 0.55 | WLOS_0017_tumor |
| | 0.54 | 0.57 | 0.53 | 0.57 | 0.55 | 0.58 | 0.95 | 1.00 | 0.56 | 0.59 | 0.55 | 0.59 | WLOS_0017_Plasma_1 |
| | 0.58 | 0.54 | 0.57 | 0.53 | 0.58 | 0.54 | 0.60 | 0.56 | 1.00 | 0.92 | 0.64 | 0.59 | WLOS_0020_tumor |
| | 0.52 | 0.57 | 0.51 | 0.56 | 0.53 | 0.57 | 0.55 | 0.59 | 0.92 | 1.00 | 0.58 | 0.63 | WLOS_0020_Plasma_1 |
| | 0.58 | 0.54 | 0.58 | 0.54 | 0.59 | 0.54 | 0.59 | 0.55 | 0.64 | 0.58 | 1.00 | 0.95 | WLOS_0022_tumor |
| | 0.54 | 0.58 | 0.54 | 0.58 | 0.54 | 0.58 | 0.55 | 0.59 | 0.59 | 0.63 | 0.95 | 1.00 | WLOS_0022_Plasma_1 |

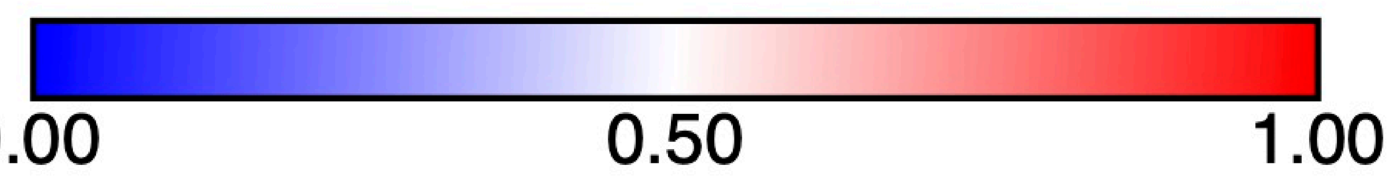

### Figure SBREEDMETRICS

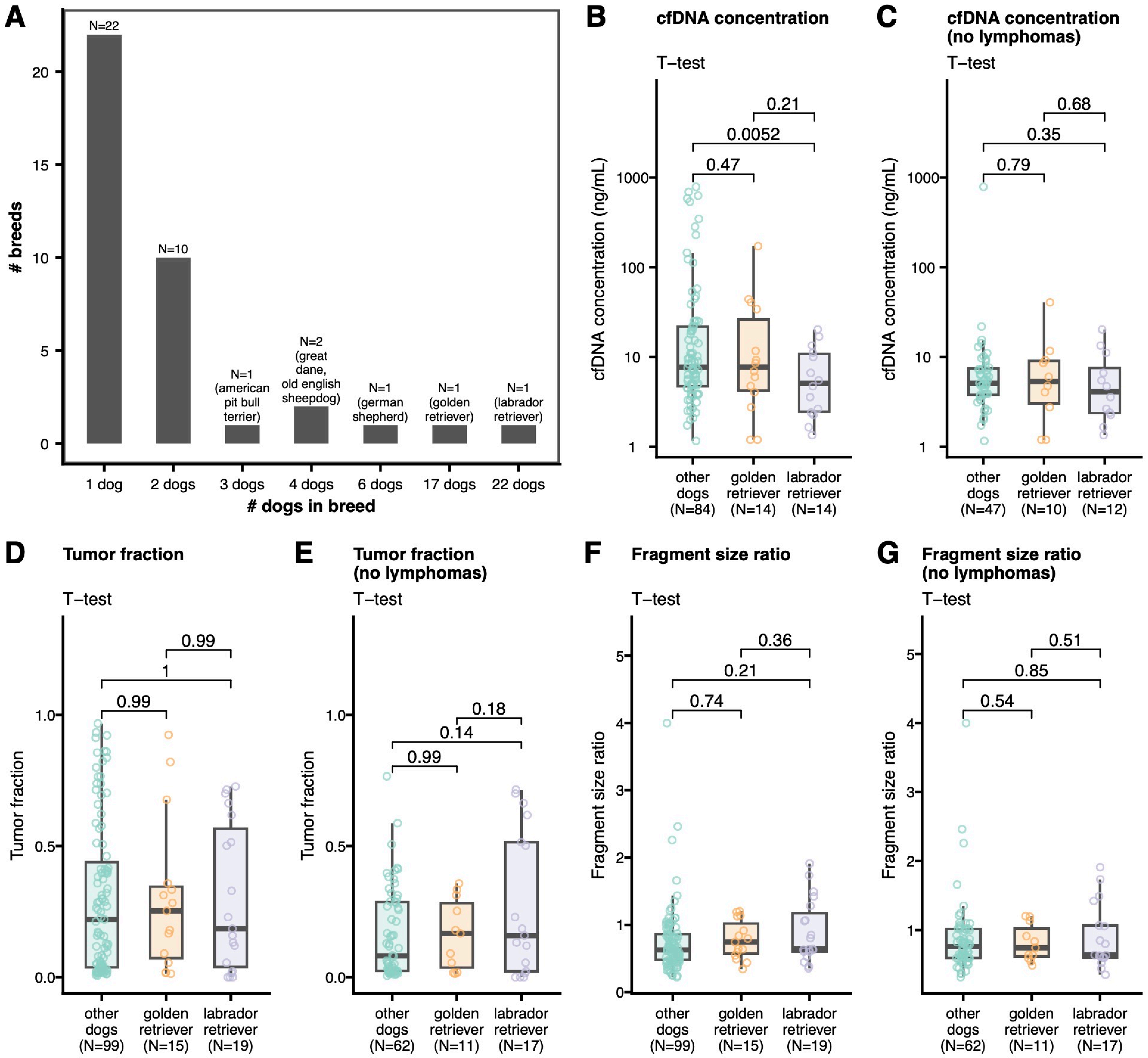

### Figure STUBES

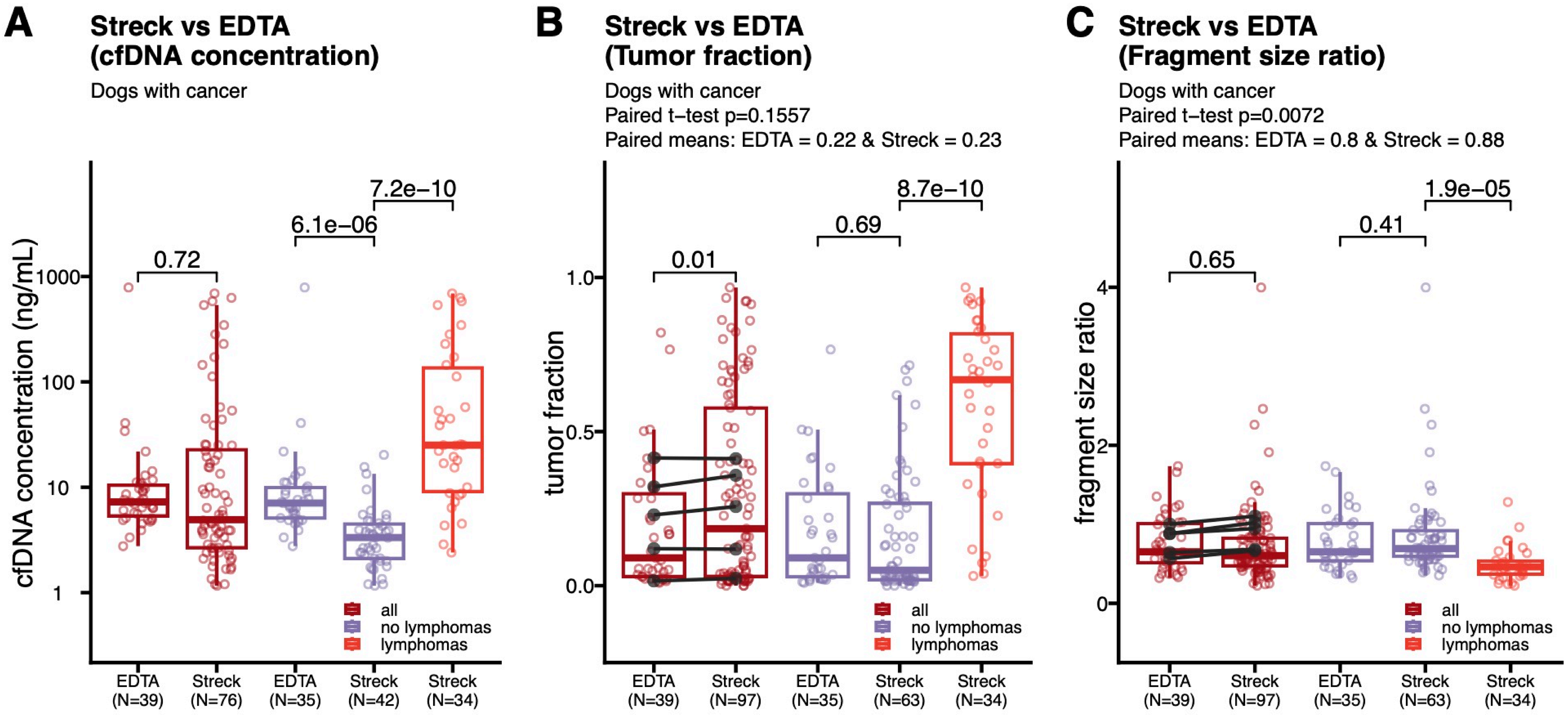

### Figure SDRAW

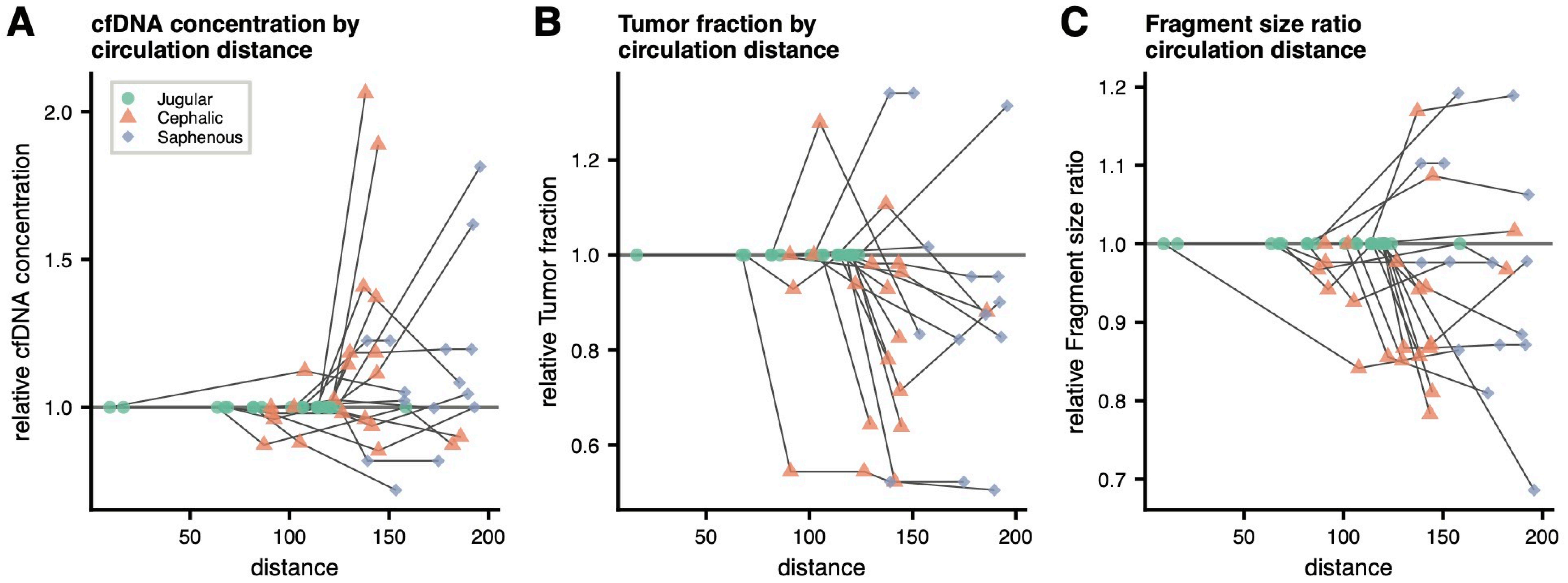

#### Figure STIME

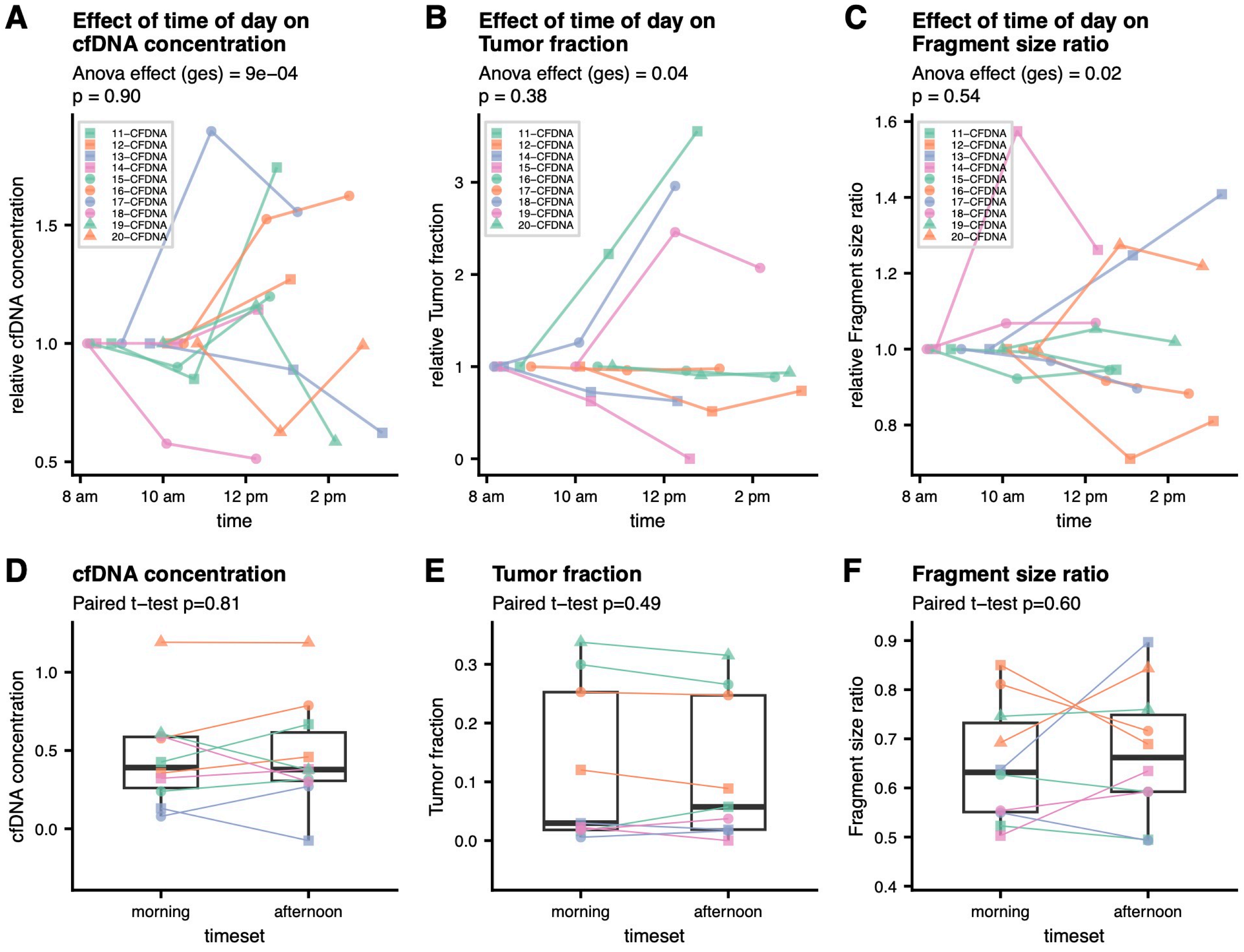

### Figure SSHORT

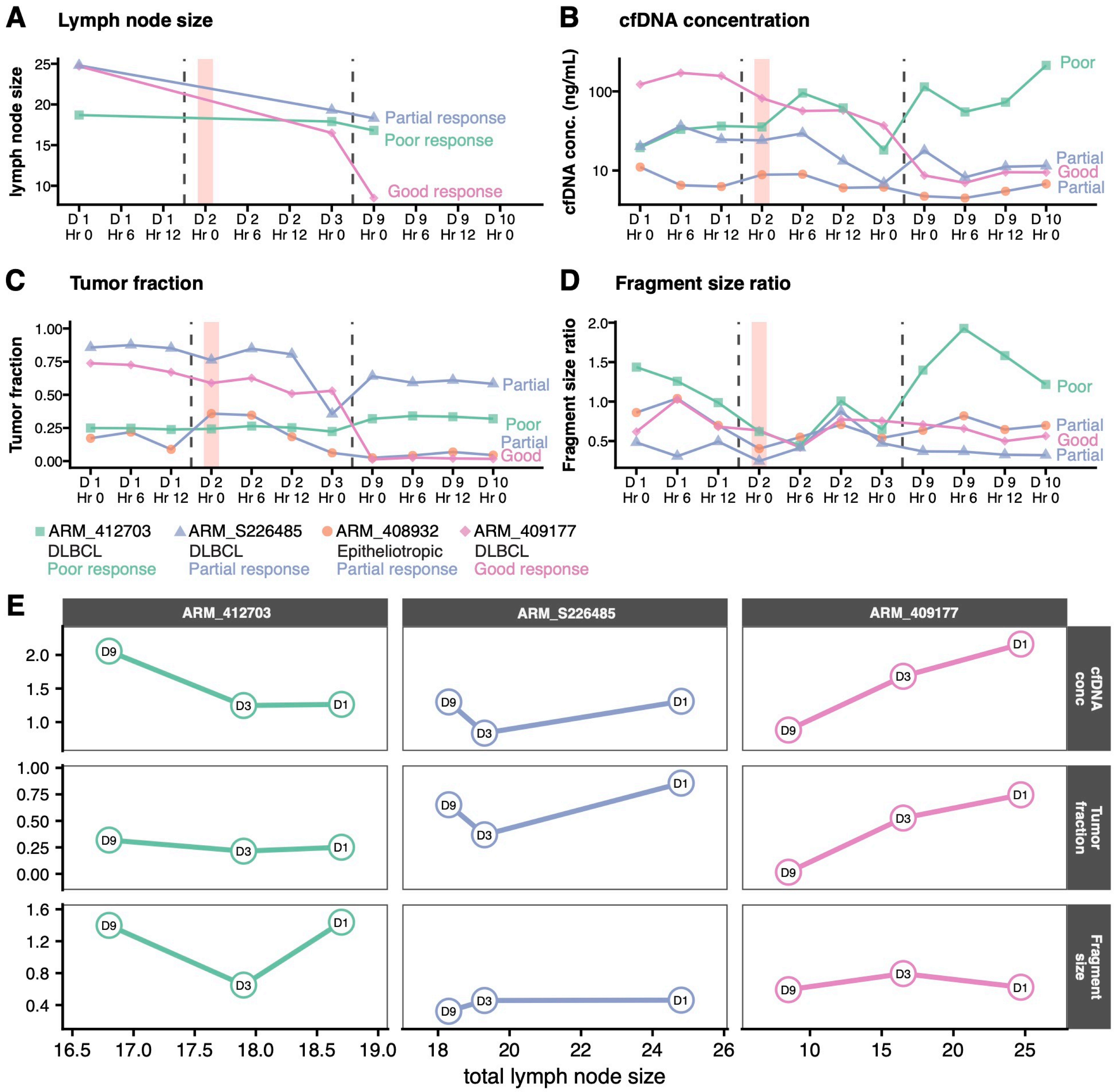

### Figure SLONBOX

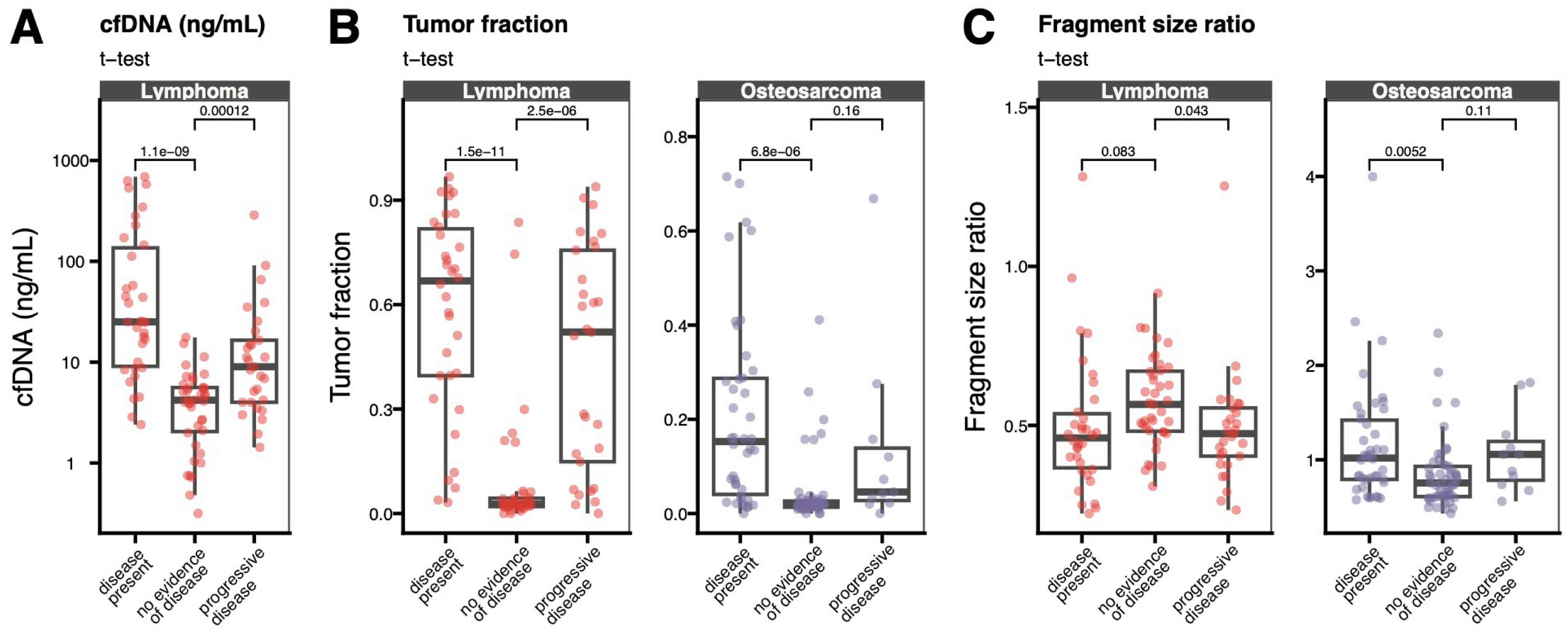

### Figure SLONGDOG_LSA_CONC

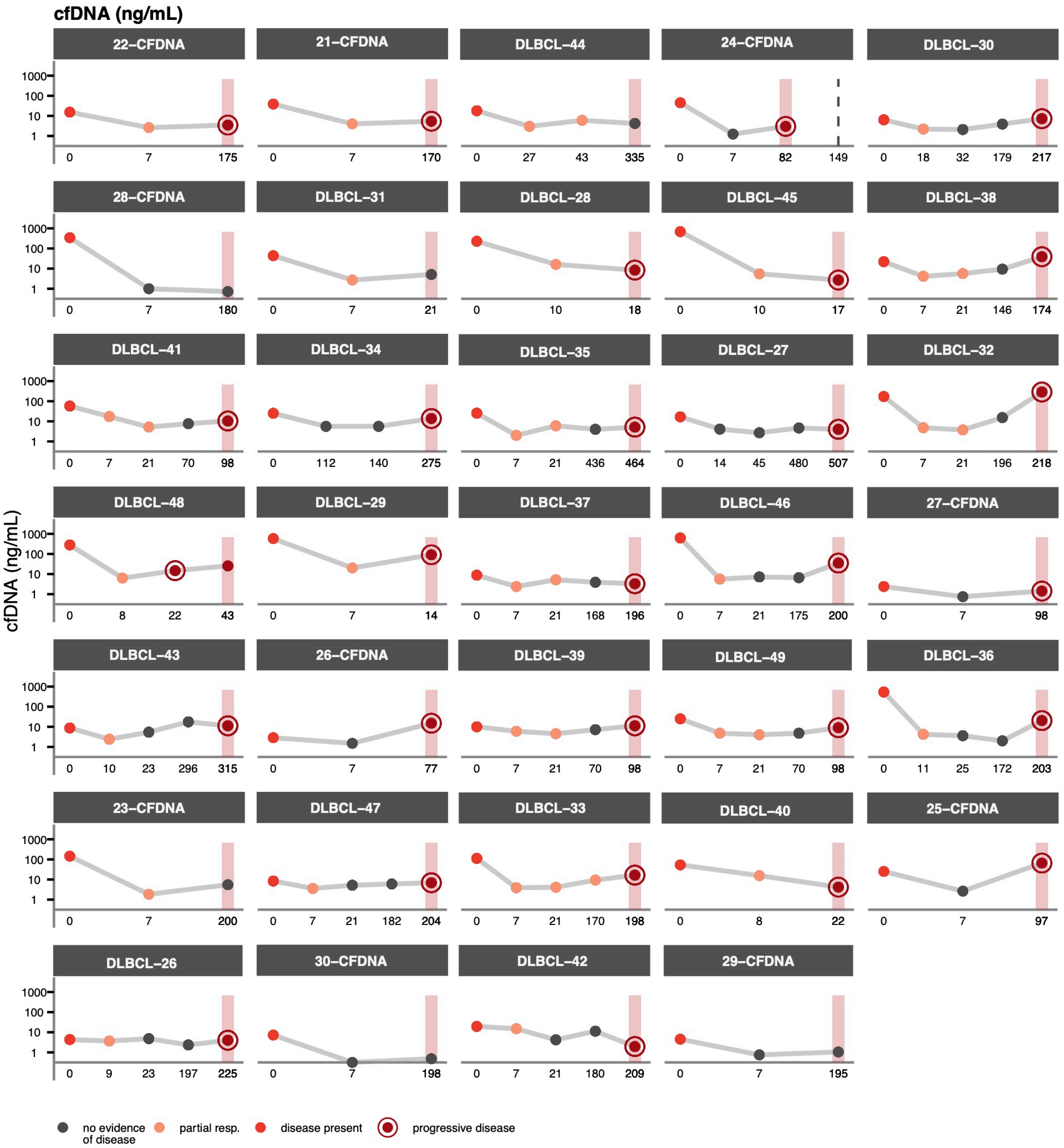

### Figure SLONGDOG_LSA_TF

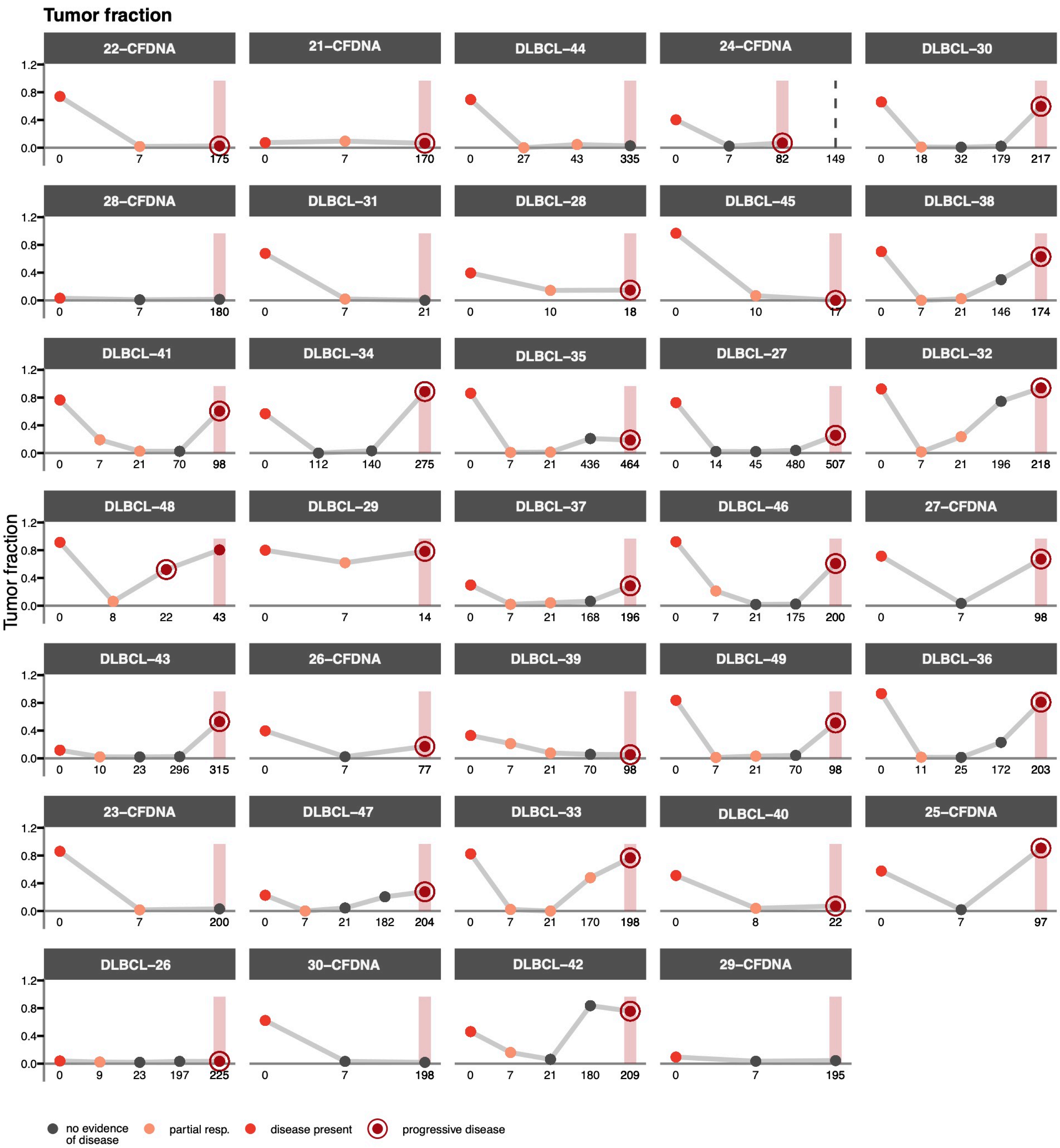

### Figure SLONGDOG_LSA_FRAG

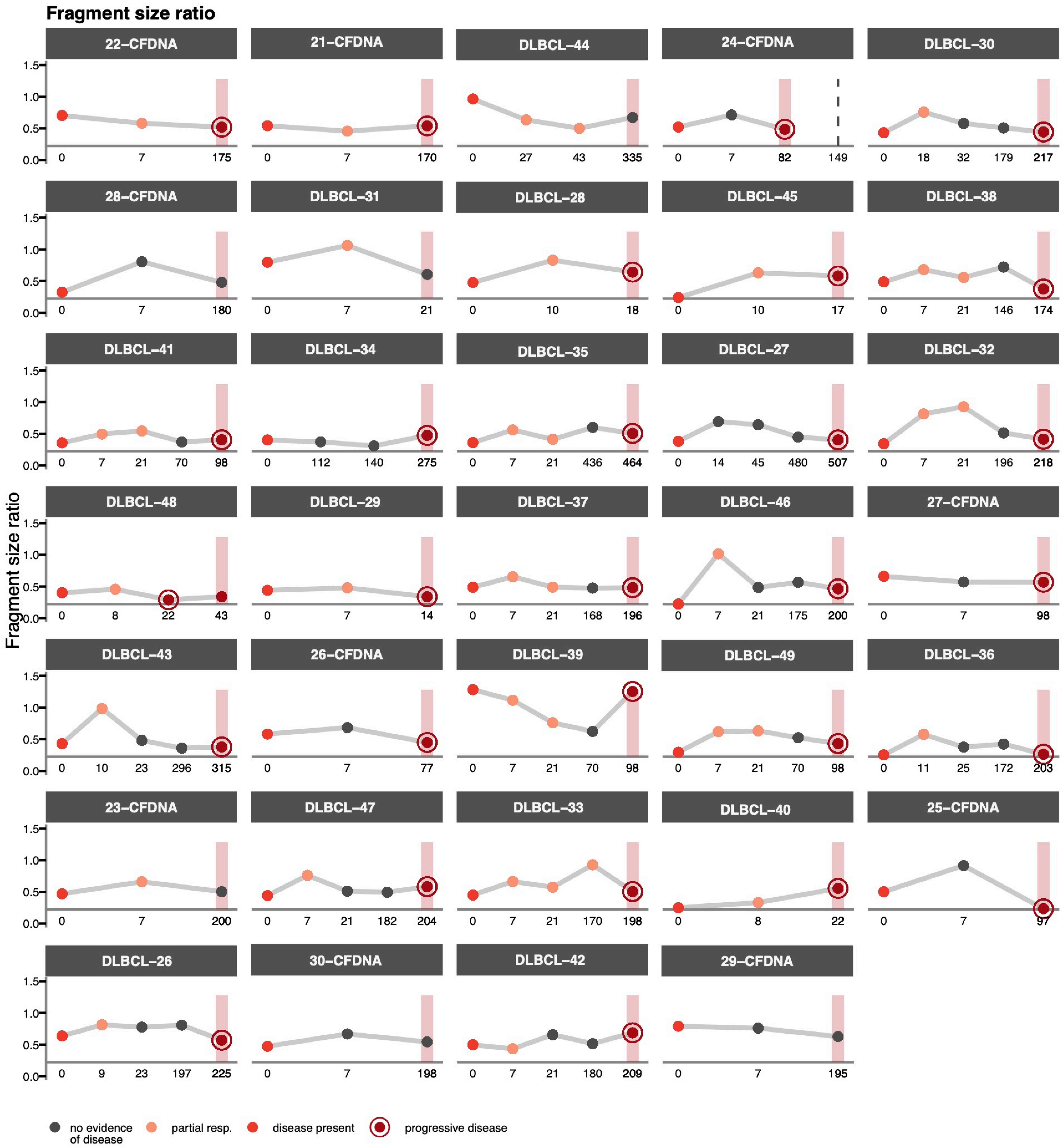

### Figure SLONGDOG_OS_TF

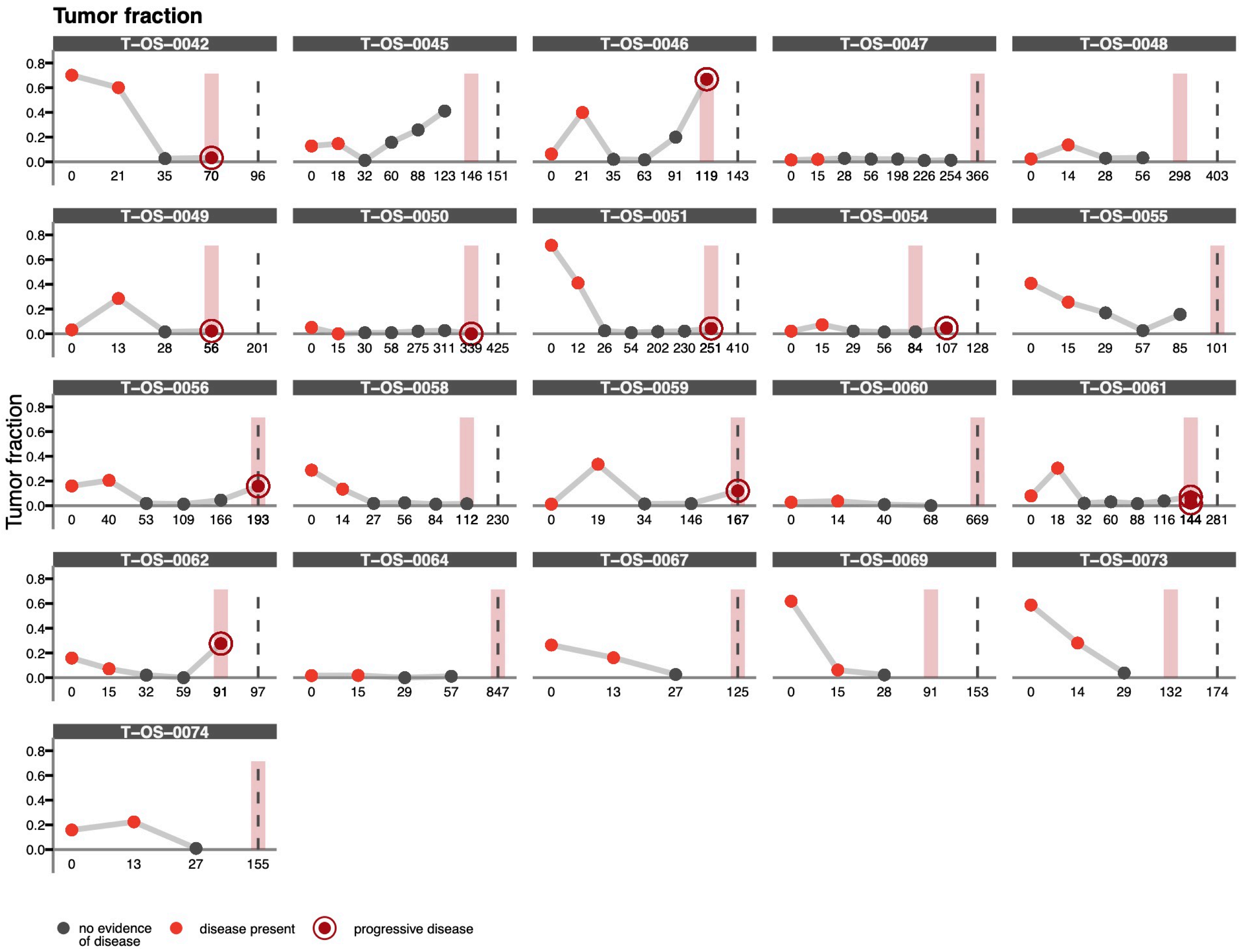

### Figure SLONGDOG_OS_FRAG

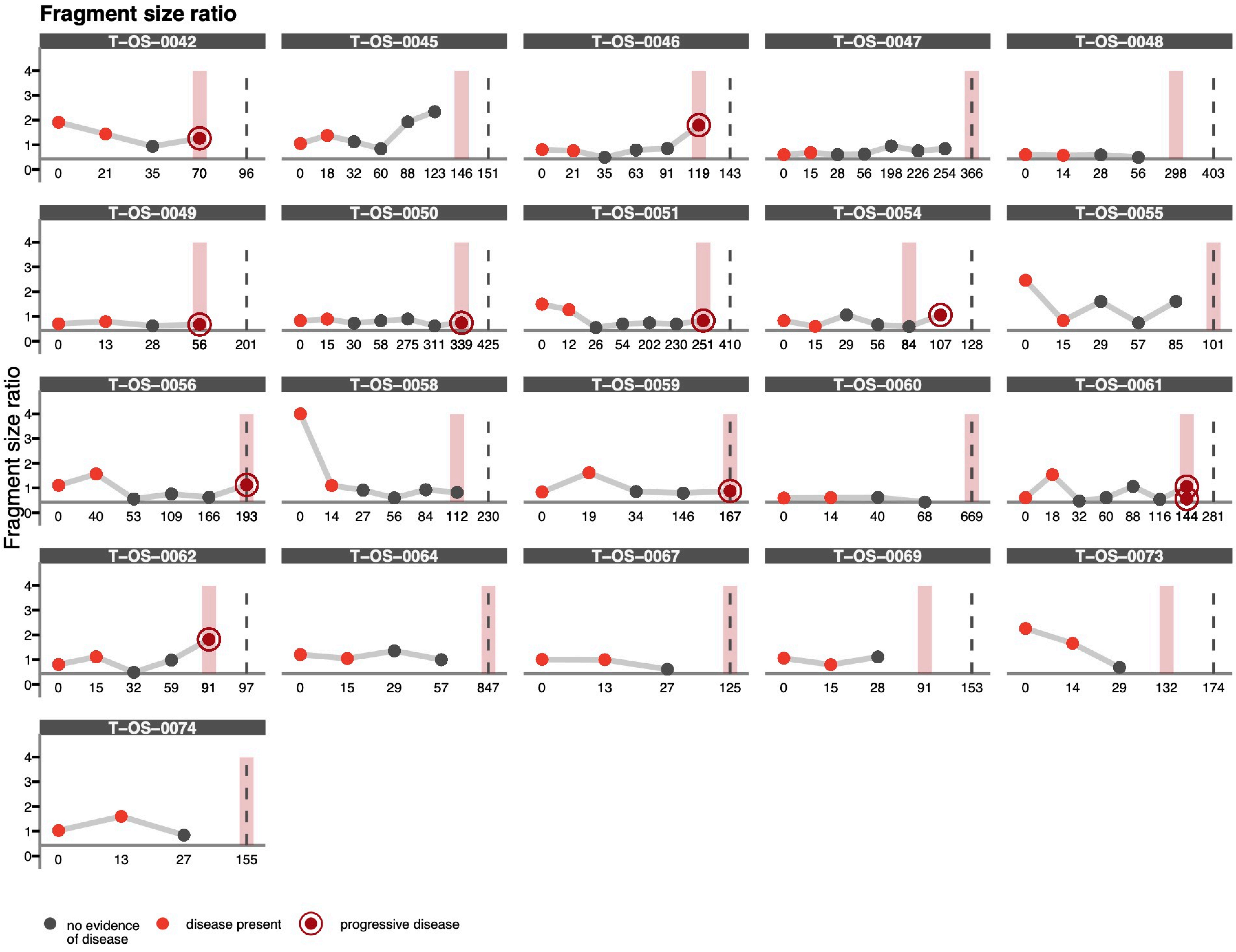

### Figure SFOREST

**A**

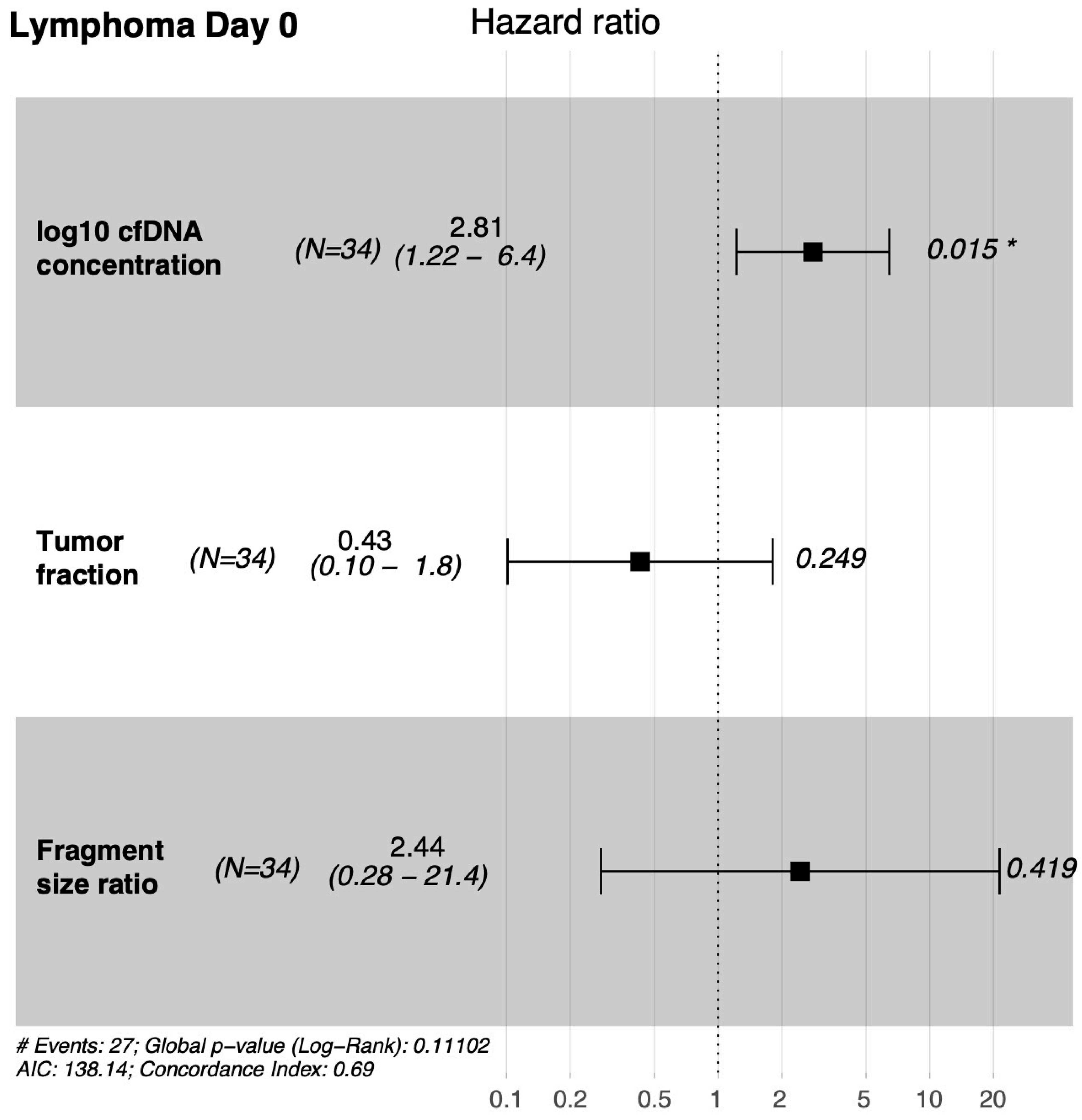


**B**

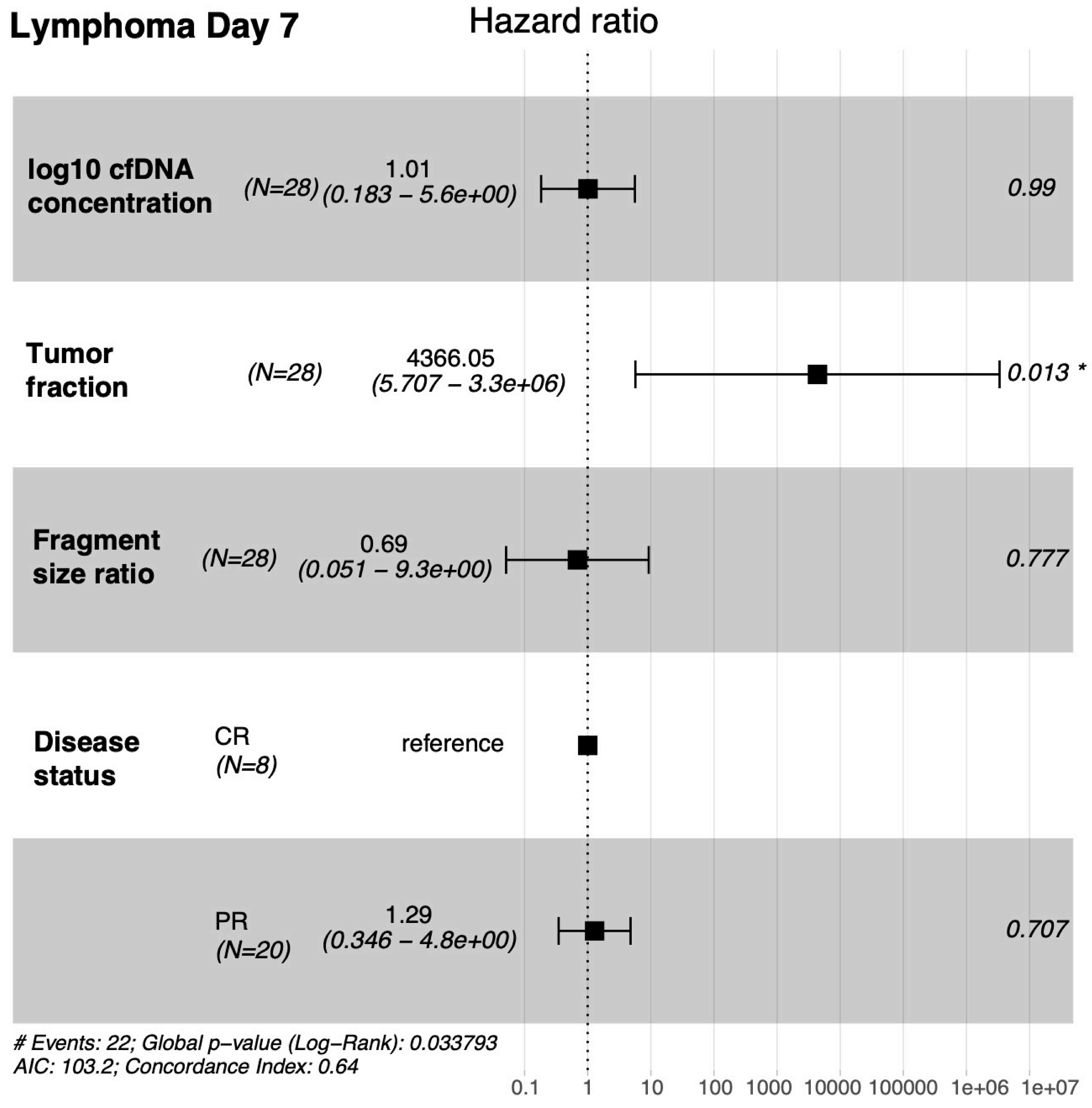
